# A flexible and versatile system for multicolor fiber photometry and optogenetic manipulation

**DOI:** 10.1101/2022.03.16.484590

**Authors:** Andrey Formozov, Alexander Dieter, J. Simon Wiegert

**Affiliations:** Research Group Synaptic Wiring and Information Processing, Center for Molecular Neurobiology Hamburg, University Medical Center Hamburg-Eppendorf, Germany

**Author notes:** these authors contributed equally to this work (Andrey Formozov, Alexander Dieter).

## Abstract

Fiber photometry is a technique of growing popularity in neuroscientific research. It is widely used to infer brain activity by recording calcium dynamics in genetically defined populations of neurons. Aside from the wide variety of calcium indicators, other genetically encoded biosensors have recently been engineered to measure membrane potential, neurotransmitter release, pH, or various cellular metabolites, such as ATP or cAMP. Due to the spectral characteristics of these molecular tools, different assemblies of optical hardware are usually needed to reveal the full potential of different biosensors. In addition, the combination of multiple biosensors in one experiment often requires the investment in more complex equipment, which limits the flexibility of the experimental design. Such constraints often hamper a straightforward implementation of new molecular tools, evaluation of their performance *in vivo*, and design of new experimental paradigms - especially if the financial budget is a limiting factor. Here, we propose a novel approach for fiber photometry recordings, based on a multimode optical fused-fiber coupler (FFC) for both light delivery and collection. Recordings can readily be combined with optogenetic manipulations in a single device without the requirement for dichroic beam-splitters. In combination with a multi-color light source and appropriate emission filters, our approach offers remarkable flexibility in experimental design and facilitates the implication of new molecular tools *in vivo* at minimal cost. The ease of assembly, operation, characterization, and customization of this platform holds the potential to foster the development of experimental strategies for multicolor fused fiber photometry (FFP) combined with optogenetics far beyond its current state.

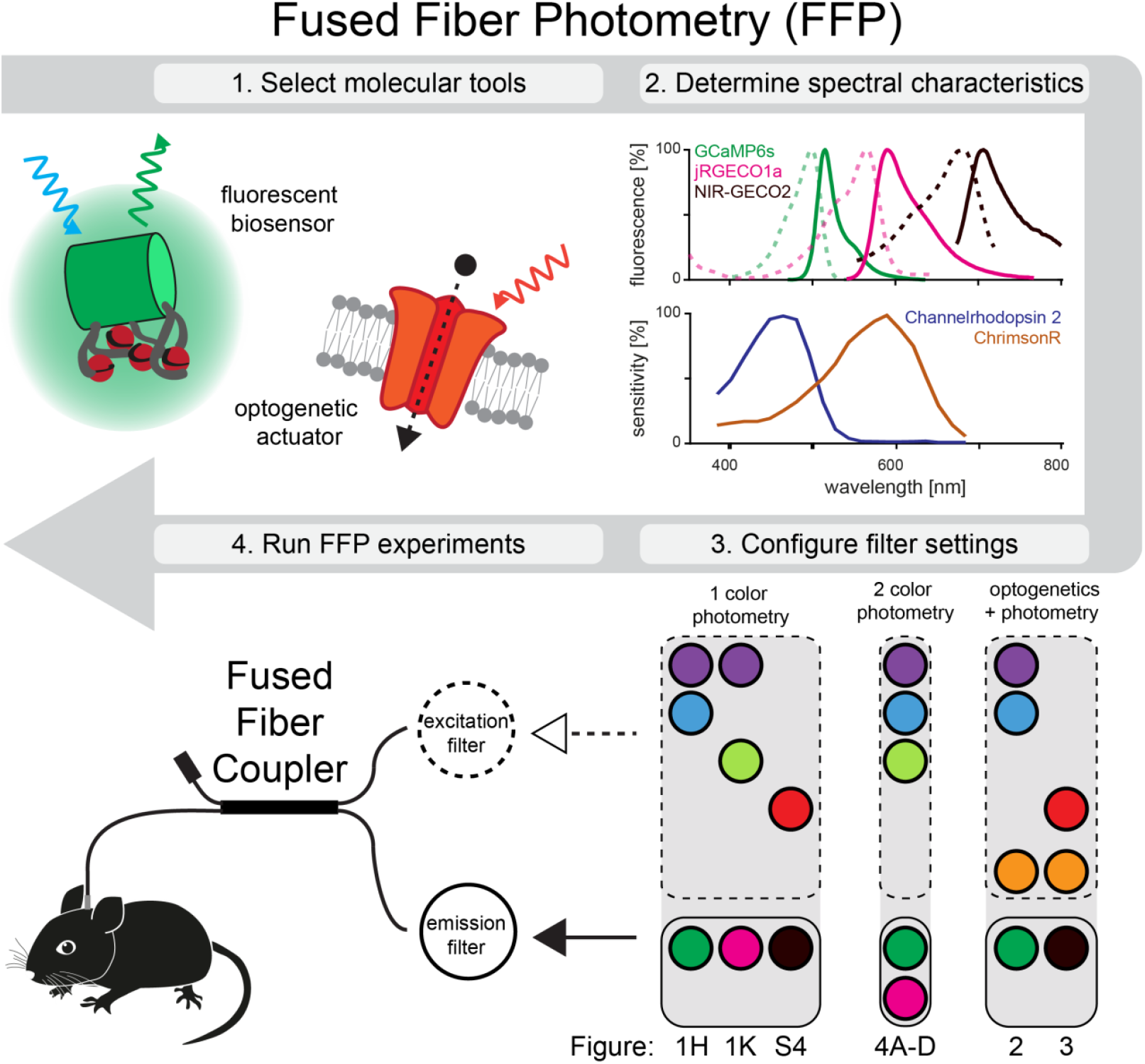

**Highlights:**

- Fused Fiber Photometry (FFP) enables straightforward monitoring and manipulation of brain activity
- FFP allows monitoring of indicators with virtually any spectral characteristics
- FFP is compatible with simultaneous optogenetic manipulation
- Easy assembly, characterization, and customization

## Introduction

Measuring neuronal activity with cellular specificity to understand brain function can be achieved both by electrophysiological and by optical methods. While electrophysiological methods offer high temporal resolution, allowing for direct measurements of neuronal spiking, optical methods have the advantage of spatial information and molecular specificity. The spatial information obtained by microscopic techniques allows for chronic imaging of the same structures at cellular and sub-cellular resolution, including organelles or synapses. However, both techniques require expensive, sophisticated equipment, they generate enormous amounts of data, and analysis is complicated. In contrast, fiber photometry measures bulk activity via an optical fiber in neuronal tissue. While temporal resolution is limited - as in most state-of-the-art optical techniques - and cellular resolution is lacking, it enables inference of the activity of molecularly defined populations of neurons at very low cost and with straightforward data analysis (Adelsberger et al., 2005; Girven & Sparta, 2017; Wang et al., 2021). Further advantages are the small size of implants and the absence of a microscopy objective, allowing for deep brain measurements, implantation of several fibers for multi-site recordings (Martianova et al., 2019), and straightforward combination with optogenetic manipulations (Li et al., 2017). Finally, experiments in freely moving animals are easily implementable. Hence, photometry received increasing popularity in recent years and – despite comparatively low complexity – has led to substantial understanding of brain function (Adelsberger et al., 2005; Gunaydin et al., 2014; Lerner et al., 2015; Wang et al., 2021).

Capitalizing on the expanding molecular toolbox of genetically encoded fluorescent biosensors, the versatility and applicability of fiber photometry is constantly increasing. Most indicators so far have been developed for monitoring of intracellular calcium dynamics which is a proxy for neural activity (Baker et al., 1971). Following the development of the widely used, GFP-based calcium sensor GCaMP (Nakai et al., 2001) and its predecessors, other indicators have been engineered to cover a broad color spectrum ranging from blue (Inoue et al., 2019; Zhao et al., 2011) to near-infrared light (Qian et al., 2019). In addition, variants with different kinetics, affinity, and dynamic range (Chen et al., 2013), as well as FRET-based sensors, have been engineered (Miyawaki et al., 1997; Rose et al., 2014). A more direct way to measure neural activity is offered by genetically encoded voltage indicators, which can achieve single-action potential resolution by directly sensing neuronal membrane voltage (Jin et al., 2012; St-Pierre et al., 2014). Besides sensors for calcium and voltage, additional sensors have been engineered to reveal the release of neurotransmitters such as glutamate (Marvin et al., 2013) or γ-aminobutyric acid (GABA) (Marvin et al., 2019), as well as a wide palette of neuromodulators, including dopamine (Patriarchi et al., 2020; Sun et al., 2018), norepinephrine (Feng et al., 2019), acetylcholine (Jing et al., 2018), and serotonin (Wan et al., 2020). Furthermore, fluorescent sensors are available for a huge variety of additional molecules found in the nervous system, reaching from ATP (Arai et al., 2018) and cAMP (Harada et al., 2017) over glucose (Mita et al., 2019) and lactate (Barros et al., 2013) to phosphate (Gu et al., 2006), magnesium (Lindenburg, Vinkenborg, et al., 2013), zinc (Lindenburg, Hessels, et al., 2013), and even pH (Ermakova et al., 2018; Tantama et al., 2011). Taken together, genetically encoded fluorescent biosensors for various ligands with distinct spectral and kinetic properties offer a huge amount of flexibility for multiplexed measurements of various parameters in the living brain. On top of monitoring neuronal activity, light of a different spectrum can be delivered to the brain via the same optical fiber. These spectral channels can, for example, be used for optogenetic manipulations of neuronal activity (i.e. light-induced activation or inactivation of neurons)(Boyden et al., 2005; Yizhar et al., 2011) or infrared neural stimulation (Richter et al., 2011; Wells et al., 2005). Similar to optical indicators of neuronal activity, there is a broad spectral palette of optogenetic tools available for neuronal manipulations (Klapoetke et al., 2014; Wiegert et al., 2017). Hence, performing photometric measurements and optogenetic manipulations via a single optical fiber provides a simple and versatile method for bidirectional communication with neural tissue. In combination with anatomically and molecularly specified expression of molecular tools, these technologies hold the potential to tackle previously unaddressed questions of brain function.

While fiber photometry was originally used in combination with a fluorescent Ca^2+^ sensitive dye in a single, column-like region of the neocortex (Adelsberger et al., 2005), subsequent work has advanced the development of fiber photometry applications in various directions, especially after the introduction of genetically encoded calcium indicators for *in vivo* imaging (Doronina-Amitonova et al., 2013; Lütcke, 2010). Later, optical fiber bundles were used to acquire photometry recordings across different brain regions simultaneously, either by distributing excitation light across different fibers with a galvanometer controlled mirror and collecting the emitted light in a single detector (Guo et al., 2015), or by using multiple fibers simultaneously and collecting the emitted light from each optical fiber with a camera (Kim et al., 2016). The latter approach was further extended with chronically implantable, high-density arrays of optical fibers (Sych et al., 2019). Besides monitoring of different brain regions, anatomically intermingled but genetically distinct populations of neurons have also been independently measured by spectral separation of green and red calcium (Meng et al., 2018) and voltage indicators (Marshall et al., 2016). Another example of the refinement of fiber photometry is the detection of calcium transients in subcellular compartments, such as axonal terminals in their target region (Qin et al., 2019).

To optimally explore and integrate various combinations of the above mentioned possibilities for individual experiments, cost-efficient, open-source, and experimentally flexible systems are desired (White et al., 2019). Recent developments of open-source solutions for fiber photometry have addressed individual aspects of this matter: Control systems and corresponding software to orchestrate optical illumination and signal detection of multiple spectral channels have been developed (Akam & Walton, 2019; Owen & Kreitzer, 2019), one of them featuring an additional, easily accessible graphical user interface (GUI) written in Python (Akam & Walton, 2019). In a different approach, a customized system based on a lock-in amplifier has enabled fiber photometry recordings at much lower light levels as compared to commercial systems, reducing photobleaching of the biosensor (Simone et al., 2018). Furthermore, the combination of fiber photometry recordings with EEG recordings in a customized hybrid implant was demonstrated (Patel et al., 2020). Finally, an open-source software platform for standardized analysis of calcium activity via fiber photometry recordings has been developed (Bruno et al., 2021). While all of these efforts greatly reduce costs and increase accessibility of fiber photometry, they either comprise complicated assemblies of optical components or lack flexibility. This arises from the fact that the assembly of optical components in current photometry systems was inspired by fluorescence microscopy setups. Thus, they are typically composed of three optical elements to spectrally separate light of different wavelengths, namely an excitation and an emission light filter and a dichroic mirror (Fig. 1A). For simultaneous measurements of multiple indicators with distinct spectra, the optical assembly needs to be replaced or extended with additional dichroic mirrors and filters, drastically increasing the cost and complexity of the system. However, as photometry records bulk fluorescence and is therefore conceptually simpler than microscopy, the complexity of the setup can be reduced: Instead of using dichroic beam splitting technologies inspired by classical fluorescence microscopy, fluorescence-based photometry measurements can be established in the brain also by fiber-based approaches.

**Figure 1.**
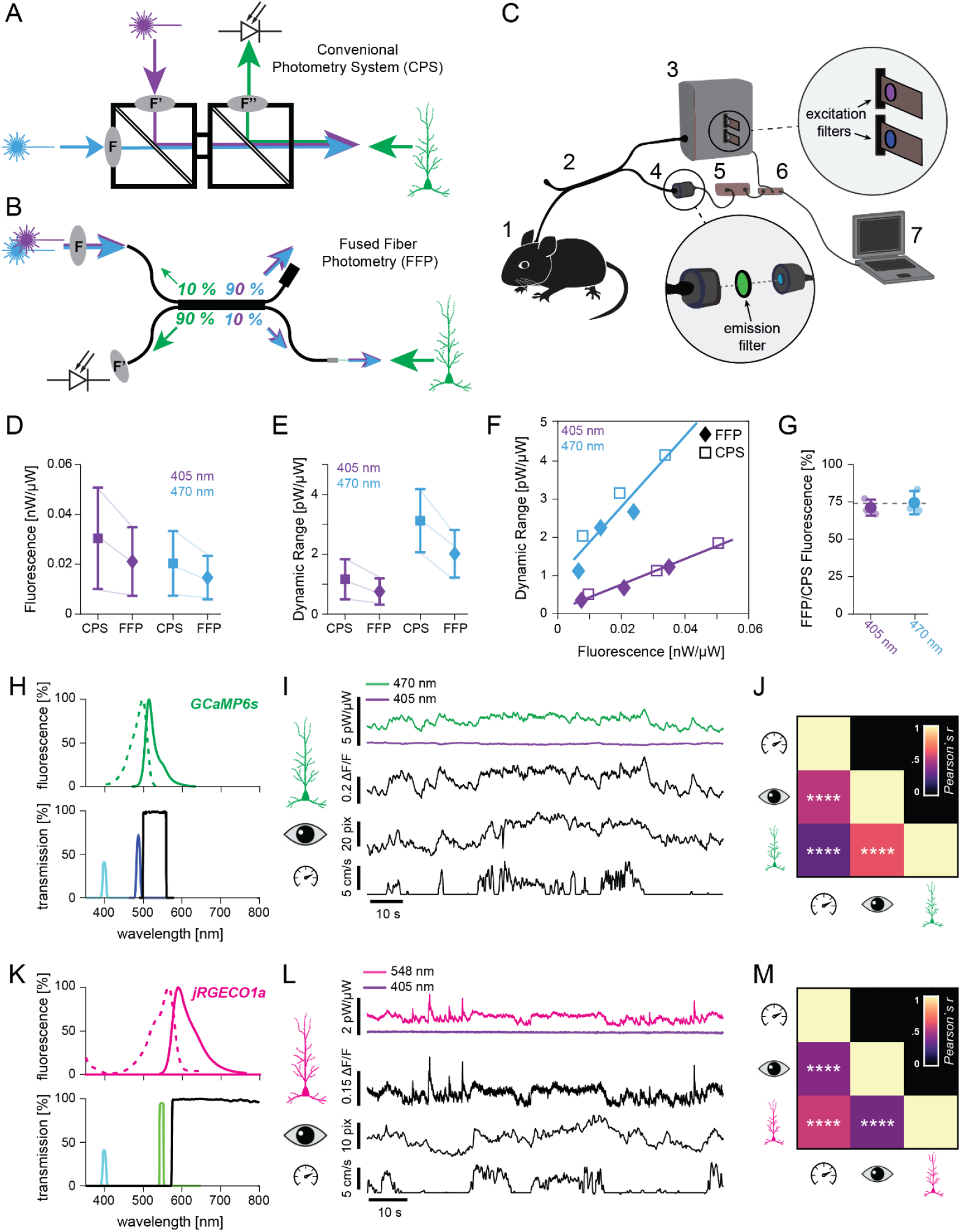
Fiber photometry recordings realized with a fused fiber photometry (FFP) system. **(A)** Scheme of a conventional photometry system (CPS) based on excitation and emission filters *F* as well as dichroic mirrors (diagonal lines). **(B)** Customized fused fiber coupler for FFP: excitation light is delivered in one branch of the coupler (top left) and split into two branches: ∼ 10 % of the power is used for the excitation of biosensors, while 90% are dissipated (top right). This configuration allows collection of 90% of emission light from the indicator by the photodetector (bottom left) through the same fiber; any backscattered excitation light is attenuated by the optical filter *F’*. **(C)** Sketch of assembled FFP system for in vivo recordings. A mouse with implanted cannula (1) is connected via the fused fiber coupler (2) to a light source (3) and a photodetector (4) with an amplifier (5). Optical filters (detailed in insets) to clean the excitation and emission light are installed in the light source and in the photodetector. The light source and photodetector are controlled by a MicroPython pyboard processor (6) connected to a computer (7) to realize indicator excitation and signal acquisition. **(D)** Within-animal comparison of median detected GCaMP6s fluorescence [nW] normalized to the excitation power [µW] recorded with CPS and FFP, when excited at 405 (purple) and 470 nm (blue). **(E)** Comparison of dynamic range, defined as the difference between the 1^st^ and 99^th^ percentiles of GCaMP6s fluorescence, normalized to the excitation power, recorded with the CPS and FFP systems. **(F)** Relationship between detected fluorescence and dynamic range of the systems. (**G)** Detected fluorescence with the FFP system, normalized to the fluorescence detected with the CPS system. Grey dashed line: Detection efficiency of the FFP system (see Figure S1) normalized to the efficiency of the CPS system. **(H)** Excitation and emission spectrum of GCaMP6f (top) and filter properties of excitation (405/10 nm, cyan; 470/10 nm, blue) and emission filters (black; bottom) to record fluorescence from hippocampal neurons expressing GCaMP6s. **(I)** Representative raw data traces of 2 min dura0074 ion: raw traces for 470 and 405 nm excitation, as well as isosbestic-corrected calcium signal (ΔF/F), pupil size, and running speed (from top to bottom). **(J)** Correlation matrix indicates positive correlations between hippocampal calcium activity, pupil size, and running. **(K)** Excitation and emission spectrum of jRGECO1a (top), properties of excitation (405/10 nm, cyan; 548/10 nm, green) and emission filters (black; bottom) to record fluorescence from hippocampal neurons expressing jRGECO1a. **(L)** Representative raw data traces of 2 min duration: raw traces for 548 and 405 nm excitation, as well as the 405 nm-corrected calcium signal (ΔF/F), pupil size, and running speed (from top to bottom). **(M)** Correlation matrix indicates positive correlations between hippocampal calcium activity, pupil size, and running speed. ****: p < 0.0001.

Here, we propose a conceptually new type of setup for fiber photometry recordings based on fused optical fibers (fused fiber photometry, FFP). The core of this approach consists of a fused multimode fiber coupler (FFC), simultaneously used for delivery of excitation and collection of emission light. This coupler can be easily combined with any TTL-controlled light source (e.g. lasers or multi-color LEDs with corresponding clean-up filters), which is available in most neuroscience labs (e.g. for optogenetic experiments or various imaging approaches) and a sensitive detector or spectrometer, resulting in a fully operational platform with manifold capabilities. In combination with a small set of optical filters, which can be flexibly exchanged in the setup, the FFP platform enables the implementation of novel indicators and actuators for fiber photometry experiments. This approach broadens the spectral bandwidth of implemented tools well beyond the possibilities of off-the-shelf setups with predefined spectral bands. The main features of this FFP platform are (1) the possibility for straightforward implementation of biosensors of any spectral range, (2) native compatibility with optogenetics via the same fiber, and (3) the potential for multi-color applications, while reducing the effort of customization to a minimum.

Our approach is currently the cheapest possible way to obtain a functional fiber photometry setup, it is highly flexible and easy to implement, even without advanced knowledge about optics. These aspects are particularly useful when entering the field of fiber photometry, pioneering experiments with various novel biosensors, or scaling up measurements, either by monitoring multiple brain regions at the same time, or by parallelizing experimental sessions with multiple animals.

## Results

### Optical properties of an FFC demonstrate the potential for FFP, promoting plug-and-play customization

FFCs are devices commonly used to split optical signals from one fiber into two fibers or, vice versa, combine signals from two fibers into one fiber. These couplers are bidirectional, and hence the same fiber port can be used for the delivery of excitation light as well as the collection of emitted fluorescence (Fig. 1B). The spitting ratio of FFCs, i.e. the ratio of light distributed between the two output fibers, is variable and depends on the length of the fused regions. In this study, we chose a customized FFC with a fiber diameter of 400 µm, a numerical aperture (NA) of 0.5, and an asymmetrical splitting ratio of 90:10 (for customizations, see supplementary table 1). We have designed an approach to implement FFCs for the recording of *in vivo* fiber photometry signals in the following way: Excitation light enters the FFC (Fig. 1B, top left), 10% of which is directed to the fiber tip with a 1.25 mm ferrule installed (brain port) in order to excite the biosensor of choice (Fig. 1B, bottom right), while 90% is guided to the dissipation port (Fig. 1B, top right). As the FFC preserves the splitting ratio in both directions, light emitted by the biosensor enters the brain port, and 90% will be guided to the detector port (Fig. 1B, bottom left), while 10% will end up at the excitation port and therefore is lost from the detected signal. In practice, additional coupling and internal losses also affect light propagation in the FFC, reducing the actual light transmittance to a range of ∼ 4-6% in the excitation path and ∼ 25-40% in the emission path, in a wavelength dependent manner (see Fig. S1 for a complete characterization). Furthermore, some fraction of excitation light also propagates back to the detector port (∼ 5 % of the power arriving at the brain port if the brain and dissipation ports are not terminated, which can be reduced to ∼1 % with termination by refractive index matching gel), requiring the installation of an emission filter in a dedicated slot of the photodetector in order to avoid contamination of the signal by excitation light (Fig. S2). As the spectrum of excitation light may not be fully monochromatic (and even relatively broad in case of LEDs), it can potentially overlap with the band of the emission filter and partially pass through it. In this case, a clean-up filter should be installed to remove any spectral component within the transmission bands of the emission filter (see Fig. S3 for more details). As modern optical filters provide light attenuation by a factor of >10^6^, this simple customization enables the direct use of the FFC for fluorescence detection in a much simpler way as compared to conventional fiber photometry systems, which are built around complicated arrangements of precisely aligned dichroic mirrors and optical filters (Fig. 1A). Hence, in combination with an appropriate excitation light source, optical filters, and a photodetector, the FFC becomes a fully functional setup for *in vivo* recordings of fiber photometry signals (Fig. 1C), which can easily be adjusted to capitalize on the whole spectrum of available GECIs. For a full list of components and pricing, see supplementary tables 2 (FFP-hardware), 3 (LED-based light sources), and 4 (optical filters used to customize the FFP system).

### Performance comparison with a conventional system

First, we assembled the FFP system in a configuration suitable to record from biosensors in the green spectrum. To this end, we installed excitation filters at 470/10 nm (to excite GCaMP in a calcium dependent manner) and 405/10 nm (for excitation at a wavelength at which GCaMP is not sensitive to calcium), in combination with a 530/55 nm emission filter to clean up the emitted fluorescence. In this configuration, we performed a side-by-side comparison against a conventional photometry system (denoted as CPS) which is based on dichroic mirrors and mainly assembled from components produced by Doric Lenses (Akam & Walton, 2019). Notably, light detection, data digitization and acquisition were performed with the same soft-and hardware, in order to directly compare the optical assembly of both approaches. With both systems, data was acquired from the same 3 mice injected with GCaMP6s under control of the CaMKII-promoter and implanted with a fiber optic cannula in the CA1 region of the dorsal hippocampus. This comparison revealed that performance of the FFP system was close to that of the CPS. Median fluorescence, normalized to excitation power, revealed approx. 30% lower signal intensities with the FFP system when compared to the conventional approach (excitation at 405 nm: 30 ± 20 vs. 21 ± 14 pW/µW for CPS vs. FFP; excitation at 470 nm: 20 ± 13 vs. 15 ± 9 pW/ µW for CPS vs. FFP; Fig. 1D). However, it is important to note that difference between animals was larger than the difference between systems. Next, we estimated the dynamic range of the setups as the difference between the 1^st^ and the 99^th^ percentiles of the acquired signal. Similar to the median fluorescence, also the dynamic range was approx. 30% lower when signals were acquired with the FFP system (405 nm: 1.2 ± 0.7 vs 0.8 ± 0.4 pW/µW for CPS vs FFP; 470 nm: 3.1 ± 1.1 vs. 2.0 ± 0.8 pW/µW for CPS vs FFP; Fig. 1E). The absolute dynamic range was linearly proportional to the mean detected fluorescence (*Pearson’s r* = 0.99/0.93, *p* = 0.0001/0.008 for excitation at 405/470 nm, respectively; Fig. 1F). As the absolute dynamic range was linearly proportional to the mean detected fluorescence (while the excitation power was kept constant), we mainly attribute the difference in performance between the FFP and the CPS to the transmission efficiency of emitted light. Indeed, we measured a transmission efficiency of 37% for the FFP system (measured at 525 nm; Fig. S1), while the transmission efficiency of the CPS system was ∼50%, i.e. transmission efficiency of the FFP system was 26% lower. This is in good agreement with the relative differences observed when comparing the FFP to the conventional system, which amounted to 29/25 % when indicators were excited at 405/470 nm (Fig. 1G). Hence, further improvements of the FFP system should aim to increase the detection efficiency.

### Recording signals of different calcium indicators *in vivo* with minimal customization

In a next step, we evaluated the biological relevance of neuronal population activity recorded with the FFP system. To this end, we virally expressed different genetically encoded calcium indicators in the CA1 region of the dorsal hippocampus, and implanted a ferrule-coupled optical fiber just above the injection site. We then obtained FFP recordings from awake, head-fixed mice on a linear treadmill, using the microcontroller-based data acquisition module of the recently developed, open-source *pyPhotometry* system (Akam & Walton, 2019). Simultaneously, we recorded locomotion and videographic images of the animals’ pupil.

Capitalizing on the flexibility of the FFP system, we first installed optical filters for photometry recordings in an animal expressing the green calcium indicator GCaMP6s (Dana et al., 2019) (as described above; Fig. 1H). Excitation of GCaMP6s was achieved with temporally interleaved light pulses at 470 and 405 nm to measure both calcium-dependent changes in fluorescence, and an isosbestic control (i.e. the wavelength of calcium-insensitive GCaMP6s excitation), allowing for the correction of baseline shifts and motion artefacts (Dana et al., 2019; Lerner et al., 2015) (Fig. 1I). In a different animal, we monitored the activity of the red calcium indicator jRGECO1a. To this end, we exchanged the filter for calcium-dependent excitation to 548/10 nm, while keeping the 405/10 nm filter to control for artefacts, as it has been shown that jRGECO1a fluorescence is minimally sensitive to calcium when excited at this wavelength (Molina et al., 2019). Emitted light was cleaned with a 575 nm long pass emission filter (Fig. 1K). In both configurations, clear calcium-modulated transients were observed when exciting the respective fluorophore at calcium-dependent wavelengths, as opposed to excitation at 405 nm, which was calcium insensitive (Fig. 1I, 1L; top).

As both arousal (inferred from pupil size) and locomotion are reportedly correlated to neural activity across a wide range of brain regions (e.g. McGinley et al., 2015), we verified functional FFP recordings by their positive correlation to the pupil size and running speed of the animal (Fig. 1J, 1M). Hence, we demonstrated that the FFP system is capable of recording biologically relevant calcium signals *in vivo*, with minimal customization enabling flexible implementation of indicators with different spectral properties. It is important to note that the choice of the excitation and emission filters is not limited to settings for detection of GCaMP and jRGECO, but can be freely determined by the experimenter. For example, settings can be adapted to allow for fluorescence recordings of cyan (CFP) or yellow (YFP) indicators, respectively. Thus, the FFP system can be natively configured for any excitation/emission combination and thus can be used as a highly customizable platform that fosters the use of the whole spectrum of available indicators.

### Combination of fused-fiber photometry and optogenetic manipulation

After demonstration of FFP recordings with a single indicator, we proceeded to combine signal collection with simultaneous optogenetic stimulation. We co-expressed GCaMP6s and the orange-light-activated cation channel ChrimsonR (Chrimson(K176R); Klapoetke et al., 2014). As both the excitation and emission spectra of the green-fluorescent GCaMP6s are well blue-shifted with respect to the activation peak of ChrimsonR (∼590 nm), we used a 575 nm long-pass filter in combination with a 585 nm LED for optogenetic stimulation. In this case, the emission filter (530/55 nm) blocks not only the light used for excitation of the indicator, but also the light used for activating ChrimsonR, while remaining residuals of activation light are subtracted from the FFP signal by the sampling algorithm (see Methods, “signal characterization”).

With this configuration, we achieved excitation of GCaMP6s and ChrimsonR via the same fused fiber coupler, using a single, multi-LED light source (Fig. 2A). We verified the absence of signal contamination by ChrimsonR activation light (585 nm) in two ways: First, we measured the spectral profile of autofluorescence of our fused fiber coupler with and without illumination from the 585 nm LED using a spectrometer (Fig. 2B). Second, we recorded traces of autofluorescence of the fused fiber coupler with the photodetector, while applying 20 stimuli of 585 light (300 μW, 1 second pulse duration, 0.1 Hz pulse frequency), both in the regular mode without amplification (as used for data acquisition throughout this study), and with a gain of 100 to amplify putative spectral cross-talk (Fig. 2C). With neither method we could detect any signal contamination, except for very mild artefacts in the photometry traces when the amplifier operated at its maximum gain of 100 (Fig. 2C, bottom; leading to an artefact of ∼0.033 pW/µW of excitation light, corresponding to ∼0.1 % of the total signal amplitude; see supplementary table 5). These results suggest the effective blocking of the activation light by the emission filter and the sampling algorithm in the practically relevant range of fluorescence intensities.

**Figure 2.**
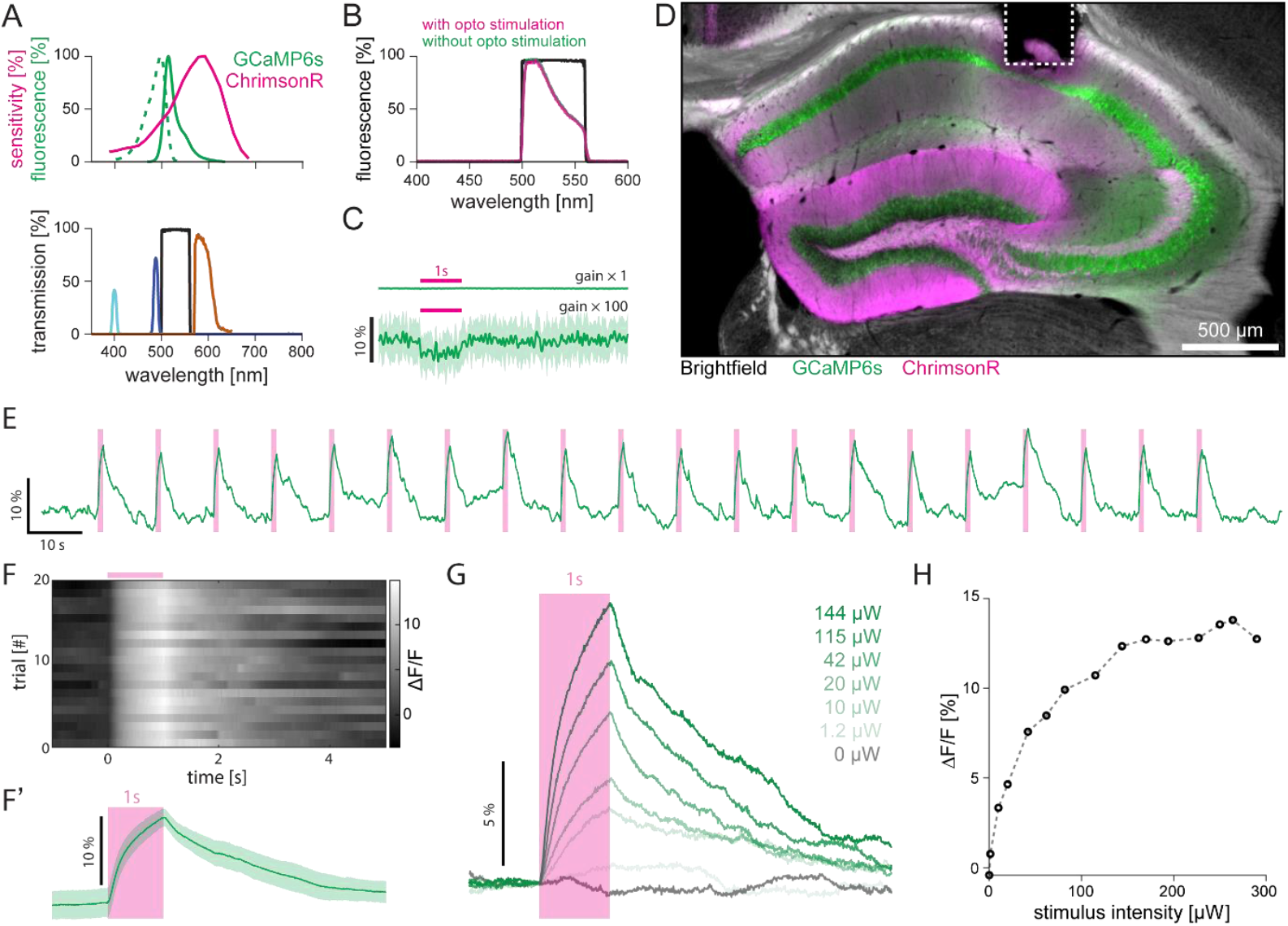
*In vivo* FFP recordings combined with optogenetic stimulation. **(A)** Excitation and emission spectrum of GCaMP6s and activation spectrum of ChrimsonR (top) along with filtered LED excitation and stimulation light (405 nm, cyan; 470 nm, blue; 575 nm, orange; bottom) and emission filter band (530/55 nm, black; bottom). **(B)** Spectrogram indicating absence of optical contamination caused by optogenetic stimulation. Transmittance of the emission filter is indicated in black, while colored lines indicate the spectrum of fiber autofluorescence with (magenta) and without simultaneous illumination with orange light for optogenetic stimulation (green). The overlapping lines indicate the absence of signal contributions originating from optogenetic stimulation as the emission filter fully blocks backscattered stimulation light. **(C)** Recording of fiber autofluorescence in response to optogenetic stimulation (300 μW for 1 second: magenta line on top). Stimulation artefacts were not detectable with a gain of 1 (top), and only mildly pronounced when the detector operated at a gain of 100 (bottom). Average (solid line) of 20 individual traces (transparent lines). **(D)** Histological verification of fiber position and expression of ChrimsonR (magenta) as well as GCaMP6s (green) in the hippocampus, overlaid with an inverted bright-field image. Dashed line: lesion caused by the optical fiber. **(E)** ΔF/F (green) in response to optogenetic stimulation (300 μW for 1 second, 0.1 Hz, 20 trials), indicated by the magenta bars. **(F)** Individual traces and **(F’)** mean ± standard deviation of ΔF/F of 20 trials of optogenetic stimulation (starting at t = 0), normalized to stimulus onset. Note the occurrence of pronounced calcium transients in every trial. **(G)** Average calcium transients in response to optogenetic stimulation with increasing stimulus intensity (20 trials per conditions). **(H)** Amplitude of average calcium traces in response to optogenetic stimulation as a function of stimulus intensity.

Next, we co-expressed GCaMP6s and ChrimsonR in the CA1 region of the dorsal hippocampus (Fig. 2D). We then acquired photometry traces from CA1 *in vivo* while activating this region with optical stimulation of ChrimsonR (300 μW, 1 second, 0.1 Hz). Optogenetic stimulation reliably evoked calcium transients in CA1, with high trial-to-trial stability (Fig. 2E, F). Modulating the radiant energy from 0-300 µW, we confirmed that the response strength scaled as a function of stimulus intensity, reaching saturation around ∼150-200 µW (Fig. 2G, H). This data demonstrates that fiber photometry recordings and optogenetic manipulations can be easily combined in a single device with the fused fiber coupler.

### Extending the FFP toolbox to the near-infrared spectrum

Recently, calcium indicators with emission spectra in the near-infrared range (NIR) have been engineered. These indicators include NIR-GECO (Qian et al., 2019, 2020) and iGECI (Shemetov et al., 2021), which are based on the fluorescent proteins mIFP and miRFP670-miRFP720, respectively. However, they have not yet found broad application, especially *in vivo*. This might result from the lack of commercial and open source hardware, and costly adaptation of set-ups to the near-infrared spectrum. In contrast, customization of the FFP system to operate with the near-infrared indicator NIR-GECO2 only requires the purchase of a single optical filter (in our case, a 700 nm long-pass filter for emission detection; Fig. S4A), which is installed in the photodetector. As the full published excitation spectrum of NIR-GECO2 is calcium sensitive and, therefore, an isosbestic point is missing, we excited NIR-GECO2 only in its most calcium sensitive range with a 650/40 nm band-pass filter, omitting a second spectral excitation band.

Switching the FFP to this configuration, we first estimated expression of NIR-GECO2 by exciting the indicator in injected mice, and compared it to the fluorescence obtained in non-injected wildtype mice: Both the absolute fluorescence (Fig. S4B) and the bleaching (Fig. S4C) were significantly higher in NIR-GECO2 injected mice (WT vs. NIR-GECO2; fluorescence: 1.21 ± 0.01 × 10^−3^ vs. 1.8 ± 0.3 × 10^−3^ nW/µW, p = 0.007; bleaching: 1.75 ± 0.23 × 10^−3^ vs. 10.7 ± 5.2 × 10^−3^ %/s, p = 0.008, Welch’s two-sample t-test, n = 3 WT, 6 NIR-GECO2 hippocampi), indicating successful expression of NIR-GECO2. We then recorded spontaneous NIR-GECO2 activity simultaneously with pupil size and locomotion, and indeed observed modulation of NIR-GECO2 fluorescence (Fig. S4D). However, as we could not include a calcium-independent control channel in this configuration, we have calculated the ΔF/F using the 10^th^ percentile of the 650 nm excited NIR-GECO2 fluorescence as baseline fluorescence F_0_, and report the fluorescence dynamics as ΔF/F_0_. As this measure of indicator activity does not correct for artefacts linked to motion and hemodynamics (which are most likely influenced by arousal, and hence linked to pupil size), we did not correlate the NIR-GECO2 traces with locomotion and pupil size, as we could not exclude signal modulation by sources different than calcium.

Instead, to circumvent the uncertainties linked to spontaneous animal behavior, we decided to record optogenetically evoked NIR-GECO2 transients. By co-expressing and optically activating ChrimsonR, we were able to lock NIR-GECO2-responses to an externally controlled event. Exploiting the flexibility of the FFP system, we combined the previously used 650/40 nm excitation and 700 nm long-pass emission filters for monitoring of NIR-GECO2 with a 585/22 nm bandpass filter for activation of ChrimsonR (Fig. 3A). Expression of NIR-GECO2 was verified post-mortem by confocal microscopy, which revealed a relatively weak fluorescence signal of NIR-GECO2 in the pyramidal cell layer of the CA1-region of the hippocampus (Fig. 3B). It is important to note that the excitation of NIR-GECO2 at our microscope was not ideal with respect to the maximum of the excitation spectrum (637 nm instead of 678 nm). Furthermore, the calcium-bound state of NIR-GECO2 is dim, and hence in fixed cells it is likely that NIR-GECO2 might not be in its bright state due to potentially elevated calcium levels prior to cell-death. In some cases, aggregates of NIR-GECO2 were also observed (Fig. S5), suggesting that the expression of NIR calcium indicators might need further optimization. Despite the seemingly low expression in fixed tissue, optogenetic stimulation resulted in robust NIR-GECO2 transients *in vivo* (Fig. 3C). While we could evoke robust transients in every trial in a NIR-GECO2 and ChrimsonR co-expressing animal (Fig. 3D-E, left), these transients were absent in an animal expressing NIR-GECO2 only (Fig. 3D-E, center). Instead, we observed a slower and longer-lasting increase in indicator fluorescence, likely reflecting photoconversion of NIR-GECO2 into a fluorescent state of higher brightness. This phenomenon has been reported before, when combining the green-light (560 nm) activated indicator jRGECO1a with an optogenetic actuator activated by shorter wavelengths (i.e. CheRiff, activated at 488 nm) (Dana et al., 2016; Nguyen et al., 2019). To estimate the role of possible optical artefacts, we also performed experiments in wildtype mice that were not injected with any indicator or actuator, but only implanted with optical fibers. We observed a minor artefact upon Chrimson-stimulation, which was one order of magnitude smaller than the responses observed in NIR-GECO2 expressing animals (Fig. 3D-E, right).

**Figure 3.**
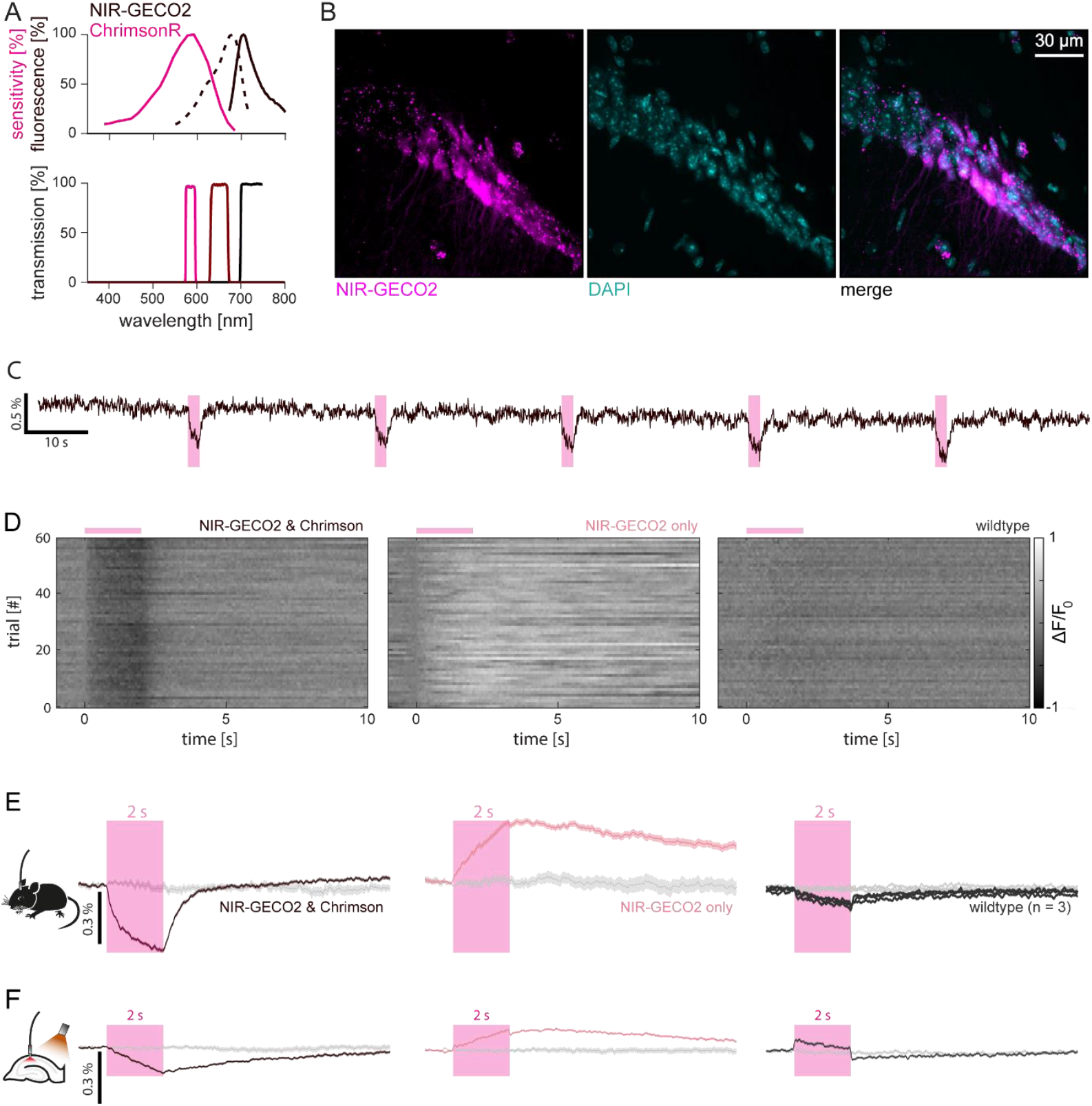
ChrimsonR-evoked calcium responses with fiber photometry recordings of NIR-GECO2. **(A)** Excitation and emission spectrum of NIR-GECO2 and activation spectrum of ChrimsonR (top) along with filter properties of excitation (585/22 nm, magenta; 650/40 nm, brown) and emission filters (700 nm long-pass filter, black; bottom)) to record NIR-GECO2 fluorescence while exciting neurons optogenetically with ChrimsonR **(B)** Histological verification of NIR-GECO2 expression (magenta) in CA1 hippocampal neurons (cyan: DAPI-staining). **(C)** Chrimson-evoked calcium transients of NIR-GECO2 (bleaching corrected 650 nm functional channel). Magenta bar: light pulse (2 s) to excite Chrimson. **(D)** Grey-scale-coded ΔF/F_0_ of 60 individual trials of stimulation with 585 nm light in an animal injected with NIR-GECO2 and ChrimsonR (left), NIR-GECO2 only (center), or without any injection (right). **(E)** Mean ± s.e.m. of the traces shown in (D). For the wildtype condition, two additional animals were measured. Grey traces indicate the mean ± s.e.m. of control trials starting at randomly selected time points of the same recording. **(F)** In vitro experiments in organotypic slice cultures verify the response of NIR-GECO2 to Chrimson-mediated excitation in the absence of motion and hemodynamic artefacts (left), the presence of photoconversion (center), and the absence of response in native slices (right). In these experiments, 595 nm stimulation light was supplied by a separate optical fiber to excite ChrimsonR in the entire slice culture.

To confirm these results and to exclude artefacts that could be caused by *in vivo* preparations (such as breathing, locomotion, or hemodynamics)(Lerner et al., 2015; Marshall et al., 2016; Zhang et al., 2021), we performed analog experiments in organotypic slice cultures. Brain slices were transduced with an identical combination of ChrimsonR and NIR-GECO2. While photometry parameters were kept constant, a fiber-coupled laser (594 nm) was used as an external light source to broadly illuminate the slice for optogenetic stimulation. As before, we observed a decrease in fluorescence when stimulating the slices transduced with NIR-GECO2 and ChrimsonR (Fig. 3F, left), while we observed a longer-lasting increase in fluorescence when stimulating NIR-GECO2 in the absence of ChrimsonR (Fig. 3F, center). In slices without any transgene expression, we observed stereotyped artefacts with a sharp on-and offset, locked to the optical stimulation (Fig. 3F, right). Although the overall magnitude of responses was smaller in organotypic slices as compared to the *in* vivo preparation, these results further verify our conclusions regarding the NIR-GECO2 applicability in combination with a red-light sensitive optogenetic actuator, and thereby demonstrate the general feasibility of FFP recordings with calcium indicators in the near-infrared range. This straightforward technical implementation within the FFP platform opens the door for future experiments employing improved versions of NIR-GECO2 or other indicators in the NIR spectrum (such as, for example, the red light activated voltage sensors of the QuasAr family (Hochbaum et al., 2014)). Taken together, the FFP system increases the flexibility of multiplexed recordings in several spectral channels or the combination of manipulation and read-out of neural activity at different wavelengths. As demonstrated here, it further enables fast and efficient benchmarking of novel fluorescent indicators in various experimental systems.

### Fused fiber photometry enables multicolor recordings with minimal customization

As demonstrated above, the combination of temporally interleaved indicator excitation and signal acquisition, together with the choice of an adequate emission filter, enables straightforward customization of the FFP system for experiments i) involving biosensors of different spectral characteristics, ii) simultaneous recording of a control channel, and iii) combination of biosensors and optogenetic actuators via the same fiber. To fully capitalize on this flexibility, we next aimed to demonstrate the suitability of the FFP system for simultaneous recordings of up to three color channels with different spectral properties showing emission peaks in the green, red, and near-infrared range. Such triple-color measurements would greatly expand the range of experimental fiber photometry applications.

To evaluate the potential and possible drawbacks of multi-color FFP, we first installed a double-bandpass emission filter (514/28 – 603/55 nm) in the photodetector, enabling simultaneous recordings of two different genetically encoded indicators in the green and in the red range (Fig. 4A). Excitation light of 470 nm (to excite the green indicator) and 548 nm (to excite the red indicator) was delivered in a temporally interleaved manner, enabling temporally – instead of spectrally – separated data acquisition of both indicators with a single photodetector (Fig. 4B). We then performed FFP recordings in organotypic hippocampal slice cultures expressing the red calcium indicator jRGECO1a and the green norepinephrine indicator grabNE1h (Feng et al., 2019). This strategy enabled us to control neuronal calcium dynamics and grabNE1h activation independently via pharmacological manipulations with bicuculline and norepinephrine, respectively (Fig. 4C). After 90 seconds of baseline recording, we applied the GABA receptor antagonist bicuculline to the slice, leading to disinhibition of the hippocampal circuit and action potential bursting (Arnold et al., 2005). A few seconds later, we observed an increase of jRGECO1a fluorescence, followed by high-amplitude oscillations of neuronal activity, indicated by the clear calcium transients detected in the red channel (Fig. 4D). We then applied norepinephrine to the slice, leading to a pronounced increase in fluorescence of grabNE1h in the green channel (Fig. 4D). While the green channel demonstrated specific responses to the application of norepinephrine, the presence of mild cross-talk in the green channel, originating from bicuculline-induced, rhythmic activity of jRGECO1a, should be noted (Fig. 4D). As the excitation spectrum of the jRGECO1a is partially overlapping with the excitation spectrum of grabNE1h, light pulses of 470 nm used for excitation of grabNE1h could to some extent also excite jRGECO1a, which still exhibits calcium sensitivity at this wavelength. To efficiently circumvent such residual cross-talk, a fast spectrometer instead of a femtowatt photoreceiver may be used for light detection in multicolor experiments. In contrast, we did not detect contamination of the red channel by the green indicator, as the 550 nm excitation light for jRGECO1a is sufficiently red-shifted with respect to the excitation spectrum of grabNE1h.

**Figure 4.**
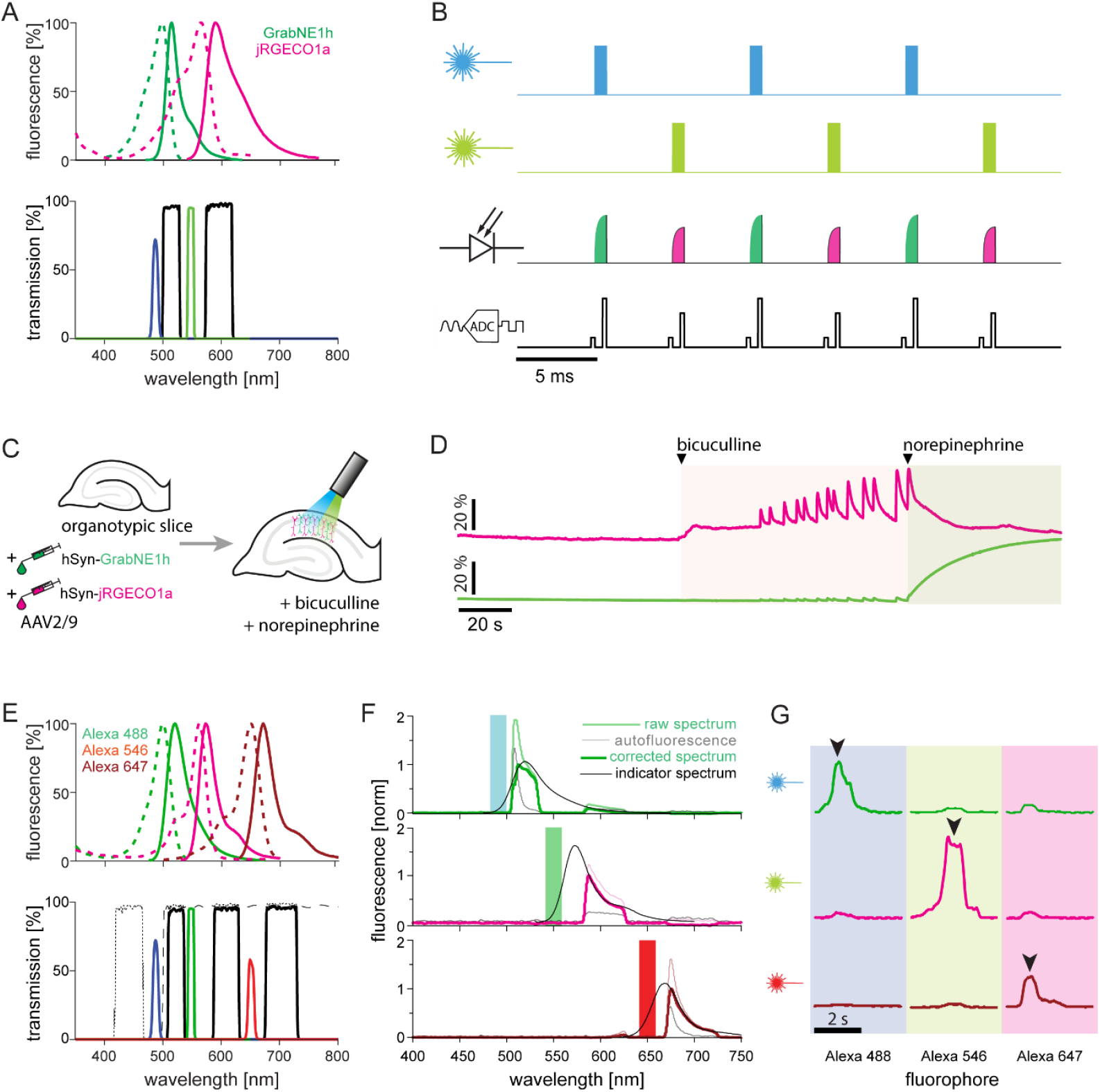
Proof of concept for multicolor FFP recordings: Experimental design and validation of temporally interleaved multi-color data acquisition. **(A)** Excitation and emission spectra of GrabNE1h and jRGECO1a (top) along with transmittance of excitation and emission filters (bottom) for the recording of two spectrally separated signals. **(B)** Sketch for the pattern of excitation and acquisition of spectrally separated indicators: blue and green light pulses are temporally interleaved for excitation of a green and a red indicator, respectively (top two traces). Fluorescence of the indicator is detected by the same photodetector (3^rd^ row from top) and digitized by an analog-digital-converter in the same, temporally interleaved manner (bottom). For each sampling point, a baseline measurement is taken before onset of the excitation light (smaller peaks in the bottom trace), which is subtracted from the measurement of the excited state (larger peaks) to correct for background illumination (e.g. room light). Sampling points of intermingled fluorescence detection can then be separated post-hoc. **(C)** Viral transduction of organotypic hippocampal slices with grabNE1h and jRGECO1a. FFP-recordings were performed in the two-color configuration, under subsequent application of bicuculline and norepinephrine. **(D)** Bicuculline-induced calcium transients (red), together with the response of grabNE1h to the application of norepinephrine. **(E)** Excitation and emission spectra of Alexa 488, Alexa 546, and Alexa 647 (top), along with transmittance of excitation and emission filters (bottom) for the recording of three spectrally separated signals. **(F)** Spectrogram of the FFP triple-color configuration (excitation light of 490, 548, and 650 nm, from top to bottom) using Alexa 488 (left), Alexa 546 (center), and Alexa 647 (right). Autofluorescence spectra (grey lines; recorded on a black sheet of paper without any indicator) have been subtracted from the raw spectra (colored thin line) to derive the actual signal of the indicator (corrected spectrum; colored line). Indicator spectra are plotted in black. **(G)** Relative intensity of temporally interleaved photometry recordings achieved with different excitation light (490, 550, and 650 nm, respectively, from top to bottom) when pointing the fiber on microscopy slides with the fluorophores Alexa 488, Alexa 546, and Alexa 647, respectively, from left to right.

In a second set of experiments, we used microscopy slides covered with the fluorophores Alexa 488, Alexa 546, and Alexa 647 (Fig. 4E, top), whose emission spectra approximately match those of GCaMP, jRGECO1a, and NIR-GECO2. To this end, we used an off-the-shelf four-bandpass emission filter (440/20 -521/21 - 607/34 - 700/45 nm) (Fig. 4E, bottom). As the photodetector used for FFP also enables stacking of different emission filters, the 440/22 nm band, which is irrelevant for this combination of fluorophores, was blocked by an additional 500 nm long pass emission filter. First, we verified these settings with recordings of the spectral characteristics of the signal obtained from different indicators. Recorded spectra closely matched reported spectra of fluorophores after the subtraction of autofluorescence background (Fig. 4F). Next, we performed temporally interleaved photometry recordings by delivering excitation light of 488/10, 548/10, and 650/10 nm, while moving the fused fiber over microscopy slides covered with the fluorophores. Each fluorophore showed a specific response to the corresponding excitation light (as indicated by the arrowheads in Fig. 4G), demonstrating that temporally intermingled multi-color recordings could achieve detection of distinct signals from all three channels. Yet, small fractions of residual autofluorescence and – as in the case of dual-color configuration – unspecific cross-talk was observed: For example, blue light may produce significant autofluorescence and can excite not only the indicator in the green spectrum, but to some lower degree also red and infrared indicators. By replacing the photodetector with a fast spectrometer, these contributions of autofluorescence could be corrected and the cross-talk could be overcome, as it permits frame-by-frame spectral fitting and extraction of signal contributions of each fluorophore individually (Meng et al., 2018). Thus, FFP opens up a perspective of temporally intermingled recordings of up to three spectral bands with a single multi-bandpass emission filter – a goal which would require much costlier and more delicate customization with classical fiber photometry setups based on optical channel separation. It should be noted that the signal quality in these experiments could likely be improved by customized filters, which will maximize the emission bands for each indicator and improve blocking efficiency. This novel recording strategy for fiber photometry systems promises an immediate impact on the set of available experimental designs for interrogation of biological circuits, as the hardware can easily be adapted to match novel indicators and actuators. Thus, the FFP system may significantly broaden the experimenter’s toolbox and facilitate the application of novel tools for *in vivo* applications.

## Discussion

In the present work, we propose a novel concept for fiber photometry recordings based on a single fused fiber coupler, suitable for the interrogation of neuronal circuits using fluorescent biosensors and optogenetic actuators. We demonstrated monitoring of calcium indicators in three different spectral ranges, including the near-infrared range, which has not been implemented in fiber photometry experiments so far. Further, we demonstrate straightforward combinations of fiber photometry with optogenetics in the same device. Similar approaches for light delivery and collection have previously been used for optical coherence tomography (Lee et al., 2013; Lorenser et al., 2013), endoscopy (Lemire-Renaud et al., 2010; Lorenser et al., 2013), temperature sensing (Musolino et al., 2016), and monitoring of opsin-eGFP expression (Diester et al., 2011), but they have not been applied to infer and manipulate neuronal activity. Notably, application of the FFP system is not limited to neuroscientific research, but it can also be employed for other excitable cells: For example, tools for simultaneous delivery and collection of light are requested for all-optical readout and control of cardiac activity (Entcheva, 2013; Entcheva & Bub, 2016; Entcheva & Kay, 2021; O’Shea et al., 2019) and skeletal muscles (Gundelach et al., 2020). In addition, the conceptual simplicity and potential for miniaturization may be of particular advantage when considering mobile devices, such as in research involving freely moving animals or even clinical applications. Furthermore, due to its flexibility and low cost, FFP might be useful for direct readout of optogenetically induced activity in the area of stimulation. For example, negative results of optogenetic modulation in combination with behavioral experiments are currently hard to interpret, as it could have technical (i.e. faulty opsin expression) or biological reasons (i.e. a modulated circuit is indeed not involved in the observed behavior). By co-injecting a calcium sensor along with the opsin and using FFP instead of optogenetic stimulation only, functional modulation of neuronal activity by the opsin can be confirmed, and negative results could be interpreted in a more conclusive way. Hence, the implementation of fused fiber couplers for optical read-out and manipulation of neuronal activity holds great potential for various applications both in basic and in applied research.

### Signal characterization

Our characterization of signals contributing to the absolute detected fluorescence suggests that the sensitivity of the FFP system is mainly limited by the autofluorescence of several components in the setup, rather than the absolute intensity of the excitation light, suggesting potential for further improvements of the system by lowering its intrinsic autofluorescence. Additional parameters affecting the signal (which we have not systematically investigated in this study) are the modal distribution of light in the fiber coupler, launch conditions of excitation and emission light, tracing through the filters and optical junctions, light propagation in the brain, reemission, and collection efficiencies. These issues were partially addressed in a quantitative manner, e.g. with measurements and simulation of light distribution and collection in the brain (Mansy et al., 2019; Pisanello et al., 2019). However, even with precise simulations of optical and technical aspects of the experiment, it is hard to estimate the impact of these factors on the final signal beyond a qualitative measure, as the biggest source of variability is most likely caused by the quality of fiber implantation, the relative position of the fiber with respect to neurons expressing the fluorescent indicator, expression levels of the fluorescent indicator and its distribution pattern.

### System customization

We demonstrated straightforward customization of the system for a range of commonly used indicators such as GCaMPs, jRGECO1a and the more recent NIR-GECO2 and GrabNE1h. It is important to underline that the flexibility of the FFP system is not limited to these configurations. Instead, indicators with any combination of excitation and emission spectra can be employed (such as CFP- and YFP-based indicators) by installing adequate optical filters. This is an important advantage that no other currently available system offers. Thus, our system fosters the use of the whole spectrum of available indicators which is otherwise restricted by the predetermined spectral configurations of available photometry setups.

### Combining fiber photometry and optogenetics

The flexibility of the FFP system enables straightforward addition of optogenetic manipulations when choosing opsins with an activation spectrum distinct to the implemented emission filter. As shown here, the red-shifted opsin ChrimsonR (Klapoetke et al., 2014) can be combined with GCaMP or even NIR-GECO2 (Qian et al., 2020). We anticipate that, exploiting the flexibility of the FFP platform, blue-light activated opsins such as ChR2 (Nagel et al., 2003), CheRiff (Hochbaum et al., 2014), or CatCh (Kleinlogel et al., 2011) can be equally easily combined with red-light emitting indicators such as jRGECO1a or jRCaMP1 (Dana et al., 2016). However, as always when combining various tools in a single preparation, some effects resulting from specific indicator and opsin properties should be taken into account: For example, when combining a blue-light gated opsin with the red calcium indicator jRGECO1a, the indicator may be photoconverted into a brighter state by optogenetic stimulation with blue light (which typically exceeds fluorescence excitation light by at least one order of magnitude) and induce changes in the baseline of the calcium indicator as was reported before (Dana et al., 2016; Nguyen et al., 2019). A similar phenomenon was found in this study, when combining the near-infrared indicator NIR-GECO2 with the red-light activated opsin ChrimsonR (Fig.4D-F). On the other hand, when pairing a green calcium indicator and a red-light sensitive opsin (e.g. GCaMP and Chrimson) one should be aware that red-shifted opsins typically show some degree of sensitivity to blue light as well, and the excitation light for the green indicator might interfere with optogenetic stimulation. However, the excitation power used for photometry recordings typically is orders of magnitude lower than the power required for optogenetic stimulation. Thus, inadvertent optogenetic stimulation is unlikely if fast-cycling opsins are used. The risk could be further reduced with the use of higher detector gains and hence lower excitation power, ultimately being limited by photon shot-noise of the detected fluorescence. In addition, the photometry light in our system is applied in very brief pulses of 0.75 ms, at maximum rates of 130 Hz, further reducing the radiant energy of blue light arriving at the neural tissue. It is important to note that many red-shifted indicators may also be excited with blue light (used to excite sensors in the green range), but the obtained fluorescence is usually dimmer than the one of the green indicators. This form of cross-talk can be determined with spectroscopic measurements of the emission of each indicator in the respective bands of a multi-bandpass optical filter.

### Multi-color applications

While we characterized the platform in the relevant spectral range of several classical indicators, any combination of excitation and emission spectra could in theory be used if the appropriate filters are installed. The actual performance of the FFP system for different single-or multi-color configurations can largely be predicted based on the spectral characteristics of the indicator and transmission/blocking characteristics of the filter (for example using the community editable fluorescent protein database FPbase (https://www.fpbase.org; Lambert, 2019)). This flexibility overcomes the hardware constraints of classical, microscopy-inspired systems based on dichroic mirrors, and hence enables the user to fully capitalize on the development of novel biosensors in spectrally distinct combinations, within or beyond the “classical” spectral properties (i.e. green and red). Furthermore, additional channels for monitoring of calcium-independent activity can easily be included (e.g. 405 nm for isosbestic excitation of GCaMPs) in order to correct for artefacts (e.g. animal motion or indicator bleaching). However, it is important to note that the bands of the emission filters determine the fraction of the emission spectrum which propagates to the detector when combining different spectral bands in one preparation, as multi-bandpass filters typically have narrower filter bands in order to accommodate additional bands of excitation light in between the emission bands (e.g. compare green bands in Fig. 1H, Fig. 4A, and Fig. 4E). Therefore, the gain of additional spectral bands comes at the cost of signal intensity of individual bands. Thus, in order to obtain the highest possible level of the signal, filters should be chosen for each application to maximize the overlap with the emission spectrum of each indicator. We note that, while we could demonstrate the general feasibility of temporally interleaved recordings at different wavelengths (Fig. 4), we have not yet succeeded in recording robust signals of two different functional indicators in mice. This was mainly due to i) the high autofluorescence of the FFC, ii) the relatively low excitation power achieved with the fused fiber coupler (due to the 90:10 splitting ratio) and iii) the relatively narrow, and non-ideal pass-bands of the off-the-shelf filters we have used in this study. Below we describe how to efficiently overcome these limitations.

### Current state and future perspectives

In the current study we propose a novel, FFC-based concept to realize fiber photometry recordings. We successfully demonstrate customization of the system to obtain recordings from indicators in three distinct spectral ranges, and combined these recordings with optogenetic manipulations. However, the signal quality obtained with the system did not yet fully reach the quality of a commercially available system based on dichroic mirrors, and the FFC system is not yet suited to realize recordings of multiple indicators with different spectral characteristics at the same time *in vivo*. While the signal quality achieved with the FFP system was sufficient for our experiments, it might reach its limit when biosensors of lower brightness are used, when expression levels are low, when expression occurs only in a small subset of neurons (due to conditional gene expression), or in small brain structures. At this point, it is important to note that the commercially available FFCs as used in this study were originally not developed with the aim to realize photometry recordings, but rather to combine light from different sources at a fixed ratio. We exploited the fact that these fused fiber couplers operate in both directions and realized photometry recordings of emitted fluorescence. Given their promise for straightforward fiber photometry experiments, their performance should be improved in the future, which can be realized in the following ways: First, current FFCs are not optimized for low autofluorescence. Thus, the fibers and epoxy used for coupler manufacturing should be chosen to have minimal intrinsic autofluorescence. Reduced fiber autofluorescence will i) result in lower background noise and hence improve the signal to noise ratio at identical excitation power and ii) enable recordings with higher excitation power, as the autofluorescence contribution to the overall signal is decreased, and consequently the photodetector has a higher dynamic range available for sensing indicator fluorescence. As the signal scales with excitation power, but the intrinsic electromagnetic noise of the photodetector does not, this aspect will further increase the signal to noise ratio of the FFP recordings. Optimizing the FFC for minimal autofluorescence will likely enable signal quality comparable to state-of-the-art commercial photometry systems. If technologically possible, Polyimide-and metal-coated fibers can be considered for FFC production as they offer minimal fluorescence (Utzinger & Richards-Kortum, 2003). Second, the efficiency of light collection from the indicator, which critically depends on the transmittance path to the detector, should be maximized. For optogenetic applications, also the transmittance of the excitation path should be further optimized, in order to achieve a higher radiant flux at the target neural population. Recently, a new type of fused fiber device, namely a wideband multimode circulator, was reported (Boudoux et al., 2019). This circulator is characterized by higher transmittance of excitation and emission light as compared to the regular fused fiber coupler, and hence promises to overcome this problem. Third, in addition to optimizing the FFC itself, additional components of the FFP setup can be improved. Rather than relying on off-the-shelf equipment, customized bandpass filters can be used in future studies in order to optimize both indicator excitation and the collection of light emitted by the indicator. Finally, a fast spectrometer could be implemented instead of a femtowatt photoreceiver in order to obtain the spectral composition of each indicator. Besides enabling multi-color photometry, spectrally resolved photometry (Meng et al., 2018), could also monitor ratiometric indicators (often consisting of a CFP-YFP pair).^1^ To this end, a 470-nm long pass filter could be used to cut excitation light for CFP, and as the spectral composition of CFP and YFP is fully captured by the spectrometer, the ratio can be determined.

In conclusion, the concept of the FFP system allows for simultaneous detection and manipulation of neuronal activity, with high flexibility at low cost, using widely-available and cost-effective components, while providing higher flexibility than all existing photometry systems. Flexibility is maximized by employing a TTL-triggered light source with a large array of spectrally distinct emitters launched into a single fiber port. Customization of the FFP system is reduced to the choice of adequate excitation and emission filters, which can be easily replaced without any previous experience in optics. This option drastically increases the degrees of freedom in the experimental design, and hence the possibilities for interrogating local neuronal populations. While the system does not yet offer the possibility for multi-color applications *in vivo*, we propose a number of modifications which should drastically improve its performance to close this gap in the near future.

## Material and Methods

### Customization of the FFC

Our device is based on a commercially available 2×2 90:10 step-index multimode fiber optic coupler (TH400R2F2B; with SMA connectors, FP400ERT fiber type, Ø400 µm core, NA 0.5; Thorlabs, Germany), for which we have requested three customized modifications (see supplementary table 1): The coupler is originally manufactured from a FP400ERT fiber which is characterized by low content of hydroxyl groups (OH). As low OH fibers are characterized by higher absorption – and presumably autofluorescence – than high OH fibers (around wavelengths of 400 nm), we have requested the coupler to be produced from a high OH fiber (FP400URT). Second, the length of 3 FFC branches (all except the brain end) was shortened to 15 cm, in order to reduce the total fiber length, and hence overall fiber autofluorescence. Third, we have requested to exchange the SMA connector on the brain end for a 1.25 mm ceramic ferrule.

### FFP system assembly

In order to turn the FFC into a functional FFP setup, it needs to be equipped with a source for excitation light and a photodetector. The excitation light was provided by a multi-color LED light source (pE-4000; CoolLED, UK), directly coupled to the SMA-connector of the FFC input port (Fig. 1B, top left). Fluorescence was detected with a photodetector system (DFD_FOA_FC; Doric Lenses, Canada) coupled to the detector end (Fig. 1B, bottom left) via a 600 µm, 0.5 NA FC-FC fiber. The 600 µm fiber becomes obsolete when replacing the SMA connector at the detector end with an FC connector, which can be directly coupled to the photodetector. Furthermore, it is recommended to terminate the “dummy end” (Fig. 1B, top right), in order to reduce backscattering of excitation light. This was achieved by installing an SMA-SMA adapter (Thorlabs, Germany) filled with refractive index matching gel (G608N3; n ∼ 1.5 at 402 nm; Thorlabs, Germany) and closed with a black dust cap (Thorlabs, Germany). For experiments, the animal is connected to the brain end of the coupler (Fig. 1B, bottom right). This connection further reduces backscattering from the brain end, as the refractive index of brain tissue has been reported to be ∼1.36 in the blue-green spectrum (Binding et al., 2011), which is close to the refractive index of the silica core of the fiber (n ∼ 1.46).

For flexible FFP recordings with different genetically encoded indicators and actuators, adequate excitation and emission filters need to be installed. Excitation filters can be easily installed, as the CoolLED (pE-4000) is already equipped with filter-holders (no filtering is needed if lasers are used for indicator excitation, as the spectrum is already narrow enough). Also, the installation of emission filters is straightforward, as the photodetector can be screwed open and is already equipped with optical lenses to ensure close to perpendicular incidence of emission light on the filter surface. These lenses leave sufficient space for accommodation of an emission filter in between. As the diameter of the filter slot is slightly larger than the diameter of standard optical filters (12.5 mm), we used a rubber tube (3 mm diameter) wrapped around the filter edges in order to properly center it on the photosensitive area of the detector (Fig. S2).

### Signal acquisition

Hardware control and data acquisition was realized with a MicroPython pyboard v1.1, with a slightly customized version of the PyPhotometry software uploaded (Akam & Walton, 2019). For control of spectrally separate channels (i.e. multi-color photometry, or single-color photometry with isosbestic control), the “single channel time division” mode of the PyPhotometry-software was used. This mode uses temporally intermingled excitation pulses of alternating wavelengths, and thus the detector is always reading a single channel (i.e. the currently excited channel) at a time (Fig. 4B). The customization is done to perform multi-color experiments and allows control of excitation with up to 4 light sources, and corresponding data acquisition from up to 4 channels from a single photodetector at sampling rates up to 65 Hz (modified by Maxime Maheu, PhD; found at https://github.com/SynapticWiringLab/pyPhotometry). The analog output channels of the MicroPython board were soldered to BNC connectors and connected to a CoolLED pE-4000 for the generation of excitation light. An analog input channel was connected to the photodetector for signal acquisition and a digital input channel was connected to an externally generated trigger to allow for data synchronization. To avoid soldering, a PyPhotometry board could be used as an alternative, which is already equipped with connectors for 2 analog and 2 digital channels (distributed by OpenEphys; https://open-ephys.org/pyphotometry-showcase).

### System characterization

After assembly of the FFC system, a characterization of its optical properties was performed. To this end, we have coupled the CoolLED pE-4000 to an optical fiber (200 µm diameter, 0.5 NA), and measured the output power at of all available wavelengths with a power meter (PM40; Thorlabs, Germany). We then connected this fiber to the input port of the FFC and measured the output power obtained at the brain end. The transmittance was then calculated as the ratio between the power measured at the brain end and the input power from the light delivering fiber (Fig. S1A-C). Transmittance of the emission path of the fused fiber coupler was measured in a similar way, but this time the light was delivered via a photometry patch cord (400 µm diameter, 0.5 NA) directly connected to the CoolLED pE-4000 at the FC end, and coupled into the brain end of the fused fiber coupler via a zirconia mating sleeve (SLEEVE_ZR_1.25; Doric lenses, Canada) to emulate fluorescence emitted by the biosensor, while the power of transmitted light was measured at the detector end of the fused fiber coupler (Fig. S1D-F). For the comparison of the FFP system with a conventional photometry system, we recorded data from the same 3 mice with both systems. From each mouse and in each configuration, we obtained 8 recordings of 5 minutes duration, switching between systems in a pseudorandomized manner to compensate for effects resulting from the sequence of recordings. We then omitted the first minute of each recording from further analysis, as an exponential decay of the signals (related to rapid photobleaching) was evident in the beginning of each recording. Remaining values were then averaged per system and animal. As both systems have different levels of intrinsic autofluorescence, we assessed this autofluorescence before each recording session and subtracted it from the acquired data

### Signal characterization

Next, we used a simple empirical model to quantify the individual components which compose the fluorescence measured at the photodetector (Fig. S6). Here, the absolute fluorescence (*F*) recorded at the photodetector is the sum of several components, namely sensor fluorescence (*F*_*Sensor*_, which can be further split into baseline fluorescence and ligand-modulated fluorescence), autofluorescence of the optical fiber (*F*_*Fiber*_) and brain tissue (*F*_*Tissue*_), non-blocked back-scattered photons of the excitation light (*F*_*Ex*_), as well as (potential) external background light sources in proximity to the photometry setup (*f*_*Background*_) and residual light from optogenetic stimulation (*f*_*Opto*_). Mathematically, this model is summarized by a simple relationship (see also Fig. S6A):

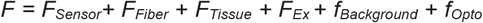

As all signal contributions (except the background light *f*_*Background*_) are proportional to the power of excitation light, it is convenient to normalize all recorded signals to excitation power, obtaining a ratio between the signal level at the detector end, and the excitation power at the fiber tip of the brain end [V/µW]. The recorded voltage can then be converted into fluorescence [nW/µW] using the characteristics of the photodetector (the conversion ratio for the photodetector used in this study is 0.76 V for 1 nW of light power at 550 nm). This conversion factor is wavelength-dependent and monotonically increases for longer wavelengths in the range 400 – 900 nm. For convenience, we have used the same conversion factor across the whole range of light used in this study. Normalized to the excitation power, these values can provide useful guidance during troubleshooting and comparisons across different experimental settings and setups. Here, we perform a detailed characterization and discuss individual signal contributions for excitation using 405/10 and 470/10 nm when recording emitted light at 530/55 nm, a setting suitable for the recording of green fluorescence, which covers the spectrum of most commonly used indicators (supplementary table 5, Fig. S6)

Optical fibers generate intrinsic autofluorescence (*F*_*Fiber*_, Fig. S6B-D). This autofluorescence depends on the material of the multimode fiber core and cladding, as well as on the distribution of guided light modes (with higher modes undergoing more absorption and reemission processes). The contribution of the autofluorescence *F*_*Fiber*_ may dominate the signal in the case of low biosensor expression levels or dim biosensors. Hence, it is generally advisable to reduce *F*_*Fiber*_ to a minimum, e.g. by photobleaching: As we demonstrated for the most commonly used wavelengths, photobleaching (using a laser light source at 405, 470, 595, and 630 nm; ∼50 mW each, for 12 hours) can reduce the autofluorescence - and hence its contribution to the absolute signal *F* - around 10-fold for excitation light of 405 and 470 nm and ∼ 2.5-fold for excitation light of 550 nm (Fig. S6B). In absolute terms, the fused fiber coupler showed remaining *F*_*Fiber*_ of ∼ 8 pW/µW, 5 pW/µW and 40 pW/µW for 405 nm, 470 nm, and 548 nm, respectively (measured with an emission filter of 530/55 nm (Edmund Optics, Stock #87-736) in the case of 405/470 nm excitation, and a green/orange double-bandpass filter (Chroma, 59012m) in the case of excitation at 550 nm. It is important to note that the commercially available fused fiber coupler used in this study is not yet fully optimized for low autofluorescence: *F*_*Fiber*_ of the FFC exceeded *F*_*Fiber*_ of an optimized low autofluorescence commercial patch cord (M127L01; Thorlabs, Germany) ∼ 21.9 and ∼ 12.2-fold for 405 and 470 nm, respectively. As the FFC does not achieve a comparable performance in terms of remaining autofluorescence, this leaves room for improvement, as also the FFC could – in principle – be manufactured from low autofluorescence materials.

Beyond the fiber autofluorescence, also the autofluorescence of biological tissue (*F*_*Tissue*_) contributes to the overall signal, in a tissue-specific manner. Here, we have measured the autofluorescence of i) paraformaldehyde-fixed brain tissue and b) live brain tissue of three non-injected, wildtype mice implanted with optical fiber cannulas into the hippocampus, in the same way as we have estimated *F*_*Fiber*_, but subtracting the fiber autofluorescence from the measured signal (Fig. S6C-D, supplementary table 5).

Another source of possible signal contamination are backscattered photons of excitation light (*F*_*Ex*_). In our system, *F*_*Ex*_ is the sum of backscattered photons from all fiber surfaces which propagate to the detector end, mainly originating from the empty dissipation (“dummy”) and brain ends of the coupler (and presumably from the coupler fusion interface). *F*_*Ex*_ amounts to 5.5 / 5 / 4.4 % of the used excitation light of 405 / 470 / 550 nm, respectively. As the backscattering at the fiber ends originates from the difference in refractive indices of the fiber and the medium surrounding it, it can be reduced by covering the dissipation end of the fused fiber coupler with refractive index matching gel, resulting in a ∼ 2-fold reduction (2.4 / 2.4 / 1.9 % for 405 / 470 / 550 nm, respectively). Additional coverage of the brain end with the gel (emulating an experimental setting in which the brain end is terminated by brain tissue) further decreased the proportion of backscattered light to 1.2 / 1.1 / 1 % for 405 / 470 / 550 nm, respectively. Finally, *F*_*Ex*_ is drastically attenuated by the use of an adequate emission filter. Modern filters guarantee an optical density (OD) > 6 defined as OD = log_10_(100% / T [%]). of backscattered excitation light is reduced > 10^6-^fold in the blocking region. In our case, this results in an overall signal contribution of 0.02 / 0.01 / 0.01 pW per µW excitation light (supplementary table 5), which is negligible in comparison to *F*_*Fiber*_ and *F*_*Tissue*_.

Ambient light sources may also induce signal contamination, which we denote as *f*_*Background*_. In the current version of the system, the data acquisition was performed with a time-binned technique developed by Akam and colleagues (Akam & Walton, 2019): This technique relies on excitation light pulses of 0.75 ms duration, flanked by brief signal acquisition periods from the photodetector. The signal preceding the excitation pulse (i.e. background illumination) is then subtracted from the signal following the excitation pulse (i.e. the desired signal plus background illumination). This way, components of *f*_*Background*_, which are usually of lower frequency than the signal acquisition, are effectively cancelled out (for a more detailed description, see Akam & Walton, 2019), and hence *f*_*Background*_ is compensated. Nonetheless, we have also assessed the background light contribution by recording signals in animals without excitation light delivered, which amounted to 5 ± 0.3 pW in our experimental setting. In contrast to other signal components, *f*_*Background*_ depicts a constant and does not scale with the excitation light. Thus, the relative contribution of *f*_*Background*_ becomes smaller with increasing excitation power in an experimental setting. Like backscattering light, *f*_*Background*_ is significantly reduced by an appropriate emission filter that fully blocks external light outside the indicator emission band.

If photometry and optogenetics are combined, the emission filter should be chosen to maximally block the light of optogenetic stimulation. Residual contamination (*f*_*Opto*_) may still contribute to the signal, inducing an artefact during illumination. However, this artefact is efficiently circumvented by the same algorithm that subtracts background light from other sources.

Finally, the main component contributing to the absolute fluorescence is the light emitted by the biosensor, *F*_*Sensor*_, which is a combination of baseline fluorescence and actual ligand-modulated fluorescence. Both components critically depend on the type of indicator and its characteristics, and are typically assessed and reported during the characterization of a novel indicator (e.g. Fig. 2 in Dana et al., 2019, for the GCaMP-family). However, it is important to note that these features are usually measured by cross-calibration of fluorescence microscopy with patch-clamp recordings in single neurons, and their extrapolation to *in vivo* photometry is not trivial, as (i) photometry signals contain activity from all neuronal compartments, and (ii) minimum fluorescence (in the absence of calcium influx) cannot be reached in the living brain. In our experiments (with mice expressing GCaMP6s under control of the CaMKII-promoter in CA1 of the hippocampus) *F*_*Sensor*_ was found to be the main - yet most variable - contributor of overall fluorescence (Fig. S6C-D, supplementary table 5). To estimate the baseline and ligand-modulated components of the fluorescence, we calculated the 1^st^ and 99^th^ percentile of signal traces, which amounted to 17 ± 12 and 12 ± 8 pW/µW (for the 1^st^ and 99^th^ percentile of signal) and 1 ± 0.15 and 0.3 ± 0.5 pW/µW (for the 1^st^ and 99^th^ percentile of dynamic range) when excited with 405 and 470 nm, respectively. Hence, only 6 and 21 % of *F*_*Sensor*_ depict actual signal modulation, which can be used to assess motion artefacts and calcium transients. The exact contribution of *F*_*Sensor*_ in different experiments depends on indicator brightness, dynamic range, expression level, and distribution in the tissue, and was found to be highly variable even when using identical virus, indicators, and brain regions, as in our experiments. Thus, *F*_*Sensor*_ may or may not exceed other background contributions in individual experiments.

### In vitro photometry recordings with organotypic hippocampal slices

Organotypic hippocampal slices were prepared from Wistar rats at post-natal day 5-7 as previously described (Gee et al., 2017). Prior to any experimental treatment, the cultures were maintained in the incubator for at least two weeks to mature at 37°C, 5% CO_2_ in a medium containing 20% heat-inactivated horse serum (Sigma H1138; supplemented with 1 mM L-glutamine, 0.00125% ascorbic acid, 0.01 mg/ml insulin, 1.44 mM CaCl2, 2 mM MgSO4 and 13 mM D-glucose) in Minimum Essential Medium Eagle (MEM; Sigma M7278). Two weeks prior to the photometry recordings, slices were transduced either with rAAV2/9-CAG-NIR-GECO2 (1×10^13^ gc/ml) and rAAV2/9-hSyn-ChrimsonR-cerulean (1.5×10^12^, in case of infrared indicator FFP configuration); or with rAAV2/9-hSyn-GrabNE1h (9.9 ×10^13^ gc/ml) and rAAV2/9-hSyn-jRGECO1a (2×10^13^ gc/ml; for dual-color experiments) by adding 0.5 µl of viral suspension on top of each slice. For FFP measurements, the brain port of the FFC was placed above the slice at the point where fluorescence was highest in order to target fluorophore-expressing neurons. In case of NIR-GECO2 experiments, an additional optical fiber (200 µm diameter, 0.22 NA) coupled to an orange laser (595 nm; Obis LS 100 mW; Coherent, Germany) was placed such that the whole slice was illuminated for optogenetic stimulation. In case of dual-color experiments, 1 µl of (+)-Bicuculline (20 mM; 0130; Tocris) and L-norepinephrine hydrochloride (10 mM; 74480; Merck Millipore, Germany) were subsequently added to the medium using a laboratory pipette.

### Multi-color application with inorganic fluorophores Alexa

Microscopy slides covered with the fluorophores Alexa 488, Alexa 546, and Alexa 647 were used as samples for the system characterization in the multi-color data acquisition mode. For this configuration, a combination of a 4-bandpass filter (440/20-521/21-607/34-700/45 nm; #87-239 Edmund optics) and 500 nm long-pass filter (#62-976, Edmund optics) was installed in the detector. To measure the spectrum of fiber autofluorescence and fluorophores, the photodetector was replaced with a spectrometer (CCS200/M; Thorlabs, Germany). As the Doric detector consists of two screwable parts, we have accommodated the emission filter into the input part of the detector, while replacing the part that contains the photodiode with an AD12F adaptor with FC connector (Thorlabs, Germany), connected to the spectrometer via an FC-SMA fiber cord (200 µm, 0.5 NA, FC-SMA; M129L01; Thorlabs, Germany). The spectrum of each fluorophore and fiber autofluorescence was then acquired during an exposure period of 5 seconds. Upon verification of the spectral components, we operated the modified PyPhotometry software in the 3-color mode, delivering sequential excitation light of 488/10 (FL488-10; Thorlabs, Germany), 548/10 (ET548/10x; Chroma, US), and 650/10 nm (FBH650-10; Thorlabs, Germany) while manually moving the 1.25 ferrule end of the coupler across microscopy slides covered with the fluorophores Alexa 488, Alexa 546, and Alexa 647.

### Animals

Experiments were performed on 11 mice (C57BL/J, *GAD2*-IRES-cre (Jackson Laboratories stock no. 028867; Taniguchi et al., 2011), and *DAT*-IRES-cre (Jackson Laboratories stock no. 006660; Bäckman et al., 2006)). Constructs were expressed independent of cre recombinase, and mice were chosen in order to minimize excess in our breeding colonies guided by the principle of the 3Rs (https://nc3rs.org.uk/who-we-are/3rs). Adult mice of either sex between 3-8 months of age were used, originating from our in-house colony (∼22°C room temperature, ∼40% relative humidity, 12/12h light-dark cycle, food and water available *ad libitum*). All experiments were performed in agreement with the German national animal care guidelines and approved by the Hamburg state authority for animal welfare (BGV, license 33/19), as well as by the animal welfare officer of the University Medical Center Hamburg-Eppendorf.

### Surgical procedures

All surgical procedures were performed under general anesthesia, either achieved by an intraperitoneal injection of a cocktail containing Midazolam, Medetomidine, and Fentanyl (MMF; 5.0, 0.5, and 0.05 mg/kg, respectively, diluted in NaCl), or by inhalation of Isoflurane (5 % for induction, ∼1.5 % for maintenance), and anesthetic depth confirmed by the absence of the hind limb withdrawal reflex. Adequate analgesia was achieved with intraperitoneal injections of Buprenorphine (0.1 mg/kg in NaCl) in the case of Isoflurane anesthesia. During anesthesia, animals were placed on a heating pad to maintain body temperature, and eye ointment (Vidisic; Bausch + Lomb, Germany) was applied to prevent drying of the eyes. The scalp of the animal was then shaved and disinfected by application of Iodide solution (Betaisodona; Mundipharma, Germany), and the animal was fixed in a stereotactic frame. To access the brain region of interest, a small incision (∼ 1 cm) was made along the scalp, and the skull was cleaned from any remaining tissue before Bregma and Lambda were stereotactically aligned. A small craniotomy (∼0.5 mm) was then performed over the region of interest (2 mm posterior and 1.5 mm lateral to Bregma to access CA1 of the hippocampus) using a dental drill. A glass pipette filled with viral suspension was then slowly lowered into the CA1 region of dorsal hippocampus (−1.6 mm relative to Bregma) using a micromanipulator. After a minute, viral suspension was slowly injected into the tissue at a speed of ∼100 nl/min by using a custom-made, manual air pressure system, until a total volume of 500-600 nl was applied. After completion of the injection, the micropipette was kept in place for at least one minute, before it was slowly retracted. To monitor calcium-modulated fluorescence across different wavelengths, we injected either rAAV2/9 encoding GCaMP6s (Chen et al., 2013) under control of the CaMKII promoter (1.25 × 10^13^ gc/ml; *AddGene* viral prep #107790-AAV9; kindly gifted by James M. Wilson), rAAV2/9 encoding jRGECO1a under control of the human synapsin promoter (4 × 10^13^ gc/ml; customized from *AddGene* plasmid #61563; kindly gifted by Douglas Kim & GENIE Project (Dana et al., 2016)), or rAAV2/9 encoding for NIR-GECO2 under control of the synthetic CAG promoter (0.5 × 10^12^ -1 × 10^13^ gc/ml; *AddGene* plasmid #159603; kindly provided by Robert Campbell (Qian et al., 2020)). For optogenetic activation, we injected either rAAV2/10 encoding ChrimsonR(K176R) and the red fluorophore tdTomato under control of the human synapsin promoter (1.13 × 10^13^ gc/ml; *AddGene* plasmid #59171; kindly provided by Edward Boyden (Klapoetke et al., 2014)) or rAAV2/9 encoding ChrimsonR(K176R) and the cyan fluorophore mCerulean under control of the human synapsin promoter (1.5 × 10^12^ gc/ml; customized from *AddGene* plasmid #59171; kindly provided by Edward Boyden (Klapoetke et al., 2014)). A detailed list of animals and viruses used in the different experiments is provided in supplementary table 6. Upon virus injection, ferrule-coupled optical fibers (2 mm length, 400 µm diameter, 0.5 NA, CFMLC15L02; Thorlabs, Germany) were slowly inserted at the same coordinates to a depth terminating 100-200 µm above the injection site and fixed to the roughened skull using cyanoacrylate glue (Pattex; Henkel, Germany) and dental cement (Super Bond C&B, Sun Medical, Japan). 5% charcoal powder was mixed with the powder of the dental cement in order to reduce optical artefacts. In addition, a head post (200-200 500 2110; Luigs-Neumann, Germany) was fixed to the skull using dental cement to allow for animal fixation during the experiments, before the incised skin was fixed to the cement with cyanoacrylate glue to close the surgical site. Once the dental cement hardened, the situs was disinfected by Iodide solution, and anesthesia was antagonized by an intraperitoneal injection of a cocktail containing Atipamezole, Flumazenil, and Buprenorphine (2.5, 0.5, and 0.1 mg/kg, respectively, diluted in NaCl) in the case of MMF-mediated anesthesia. Additional analgesia was then provided by a subcutaneous injection of Carprofen (4 mg/kg, diluted in NaCl), and Meloxicam was mixed into softened food for the three days following surgery.

### Data acquisition

Experiments were performed earliest 3 weeks after surgical procedures to allow for animal recovery, subsidence of potential inflammations, and sufficient transgene expression. Initially, animals were handled for a couple of minutes on 2-3 subsequent days to allow for habituation to the experimenter, before being taken to the setup. Subsequently, animals were habituated to head fixation (200-100 500 2100; Luigs-Neumann, Ratingen, Germany) on a linear treadmill (700-100 100 0010; Luigs-Neumann, Ratingen, Germany) for increasing amounts of time in subsequent sessions. Once animals stayed calm on the treadmill, we recorded spontaneous calcium activity, locomotion, and videography of the animal’s eye as follows: The fiber tip (brain end) of FFC was connected to the ferrule of the optical fiber implant on the animals’ head using a zirconia mating sleeve (SLEEVE_ZR_1.25; Doric lenses, Quebec, Canada) and covered with a black shrinking tube to reduce optical noise. Excitation of biosensors was achieved using a multi-LED illumination system (pE-4000; CoolLED, Andover, UK) with appropriate ⌀25 mm clean-up filters: 405/10 nm OD4 (#65-133, Edmund optics), 470/10 nm OD4 (7394, Alluxa), 548/10 nm OD6 (ET548/10x, Chroma), at intensities of 50-200 µW at the brain end. Photodetector readout and data digitization was realized with our modified version of the pyPhotometry system (Akam & Walton, 2019), using a time-binned sampling mode with baseline subtraction at a sampling rate of 65-130 Hz. In short, background illumination was measured in a time interval of 250 µs using a commercial fluorescence detector (DFD_FOA_FC; Doric lenses, Quebec, Canada). Subsequently, biosensors were excited with a light pulse of 750 µs duration, while the emitted light was measured in the last 250 µs of the excitation period. The difference between the excited state and the baseline was then taken as the intensity of light emitted by the biosensors. Mouse locomotion was measured from treadmill movement with a commercial virtual reality system, while the virtual reality displays were not operated (700-100 100 0010; Luigs-Neumann, Ratingen, Germany). Finally, the animal’s eye was monitored using a monochrome camera (DMK 33UX249; The Imaging Source, Germany) with a macro objective (TMN 1.0/50; The Imaging Source, Germany) and a 780 nm long-pass filter (FGL780; Thorlabs, Germany). Background illumination at 850 nm was provided by an infrared spotlight, while baseline pupil dilation was adjusted to be moderate using a UV-LED coupled into a polymer fiber and directed to the animal’s eye. Measurements were synchronized by custom scripts written in MATLAB (The MathWorks; Nattick, US), actuating on a NI-DAQ card (PCIe-6323; National Instruments, Austin, US).

### Data analysis

All photometry data was acquired in *.ppd format (Akam & Walton, 2019). Data analysis was performed with custom MATLAB scripts (MATLAB, The Mathworks, US). As a practically relevant measure, we used a bleaching and motion corrected signal *ΔF*/*F*, calculated the following way: First, we fitted 405 nm and calcium-dependent signals with a least-square polynomial fit of 2-5^th^ order (based on visual inspection; using MATLABs built-in “polyfit” function). We then divided signals by their respective fit, in order to correct for bleaching. Subsequently, signals were low-pass filtered (< 10 Hz; using MATLABs built-in “lowpass” function). Finally, we calculated the *ΔF*/*F* similar to Lerner et al. (Lerner et al., 2015):

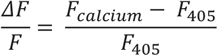

Where *F*_*calcium*_ and *F*_405_ denote the bleaching-corrected and filtered data traces of calcium-dependent and 405 nm excited indicator fluorescence. In the case of NIR-GECO2, we have calculated *ΔF*/*F*_0_, where we calculated the change in fluorescence of the active channel against its baseline fluorescence *F*_*0*_, defined as the 10^th^ percentile of the bleaching-corrected and filtered data trace:

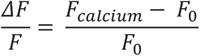

### Statistics

Correlation matrices for calcium activity, pupil size and locomotion were calculated in the following way: data traces were split in epochs of 10 s, and the Pearson correlation coefficient and its statistical significance were calculated for each epoch. Median values of all epochs for Pearson correlation and p-values were taken as a proxy for overall correlation of the signals. Welch’s two-sample t-test was used for the comparison of the signal fluorescence levels and bleaching. Everywhere throughout the manuscript, we used the following conventions for statistical significance: *: p < 0.05, **: p < 0.01, ***: p < 0.001, ****: p < 0.0001.

## Code and data availability

All code and data generated to establish the method will be made publicly available upon publication of the manuscript

## Acknowledgements

This work was funded by the Deutsche Forschungsgemeinschaft (DFG, German Research Foundation) - 178316478 - B8 to JSW. AF is supported by the Alexander von Humboldt Foundation (AvH Research Fellowship). We thank Dr. Maxime Maheu for adaptation of the PyPhotometry code to accommodate 4 separate stimulation channels. We thank Adrianna Nozownik for preparation of organotypic hippocampal slice cultures. The authors gratefully acknowledge Kathrin Sauter for plasmids preparation, Dr. Ingke Braren for virus production, and Stefan Schillemeit for technical support with histological work. We thank Prof. Dr. Thomas Oertner for providing us with the rAAV2/9-hSyn-ChrimsonR(K176R)-mCerulean construct.

## Supplementary Material

**Supplementary table 1.**
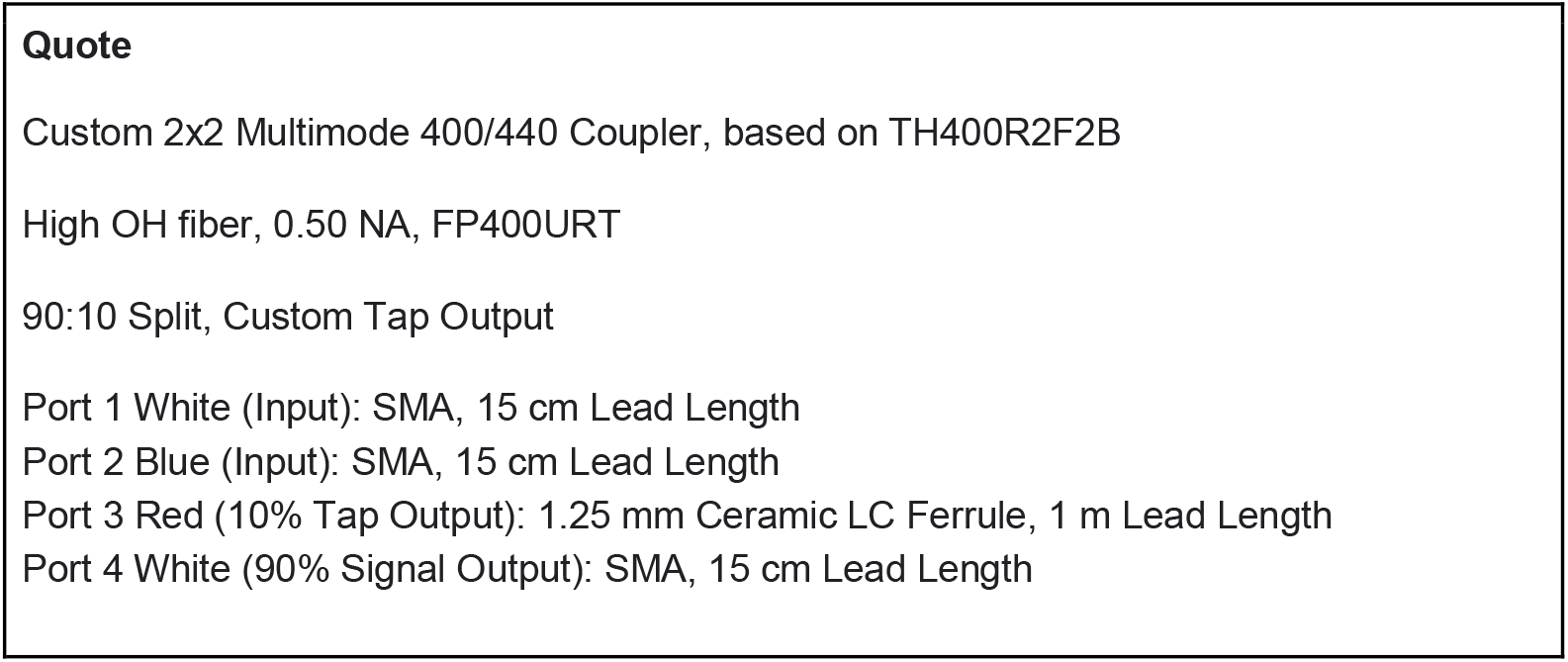
Customization parameters for fused fiber coupler. (TH400R2F2B; Thorlabs, Germany).

**Supplementary table 2.**
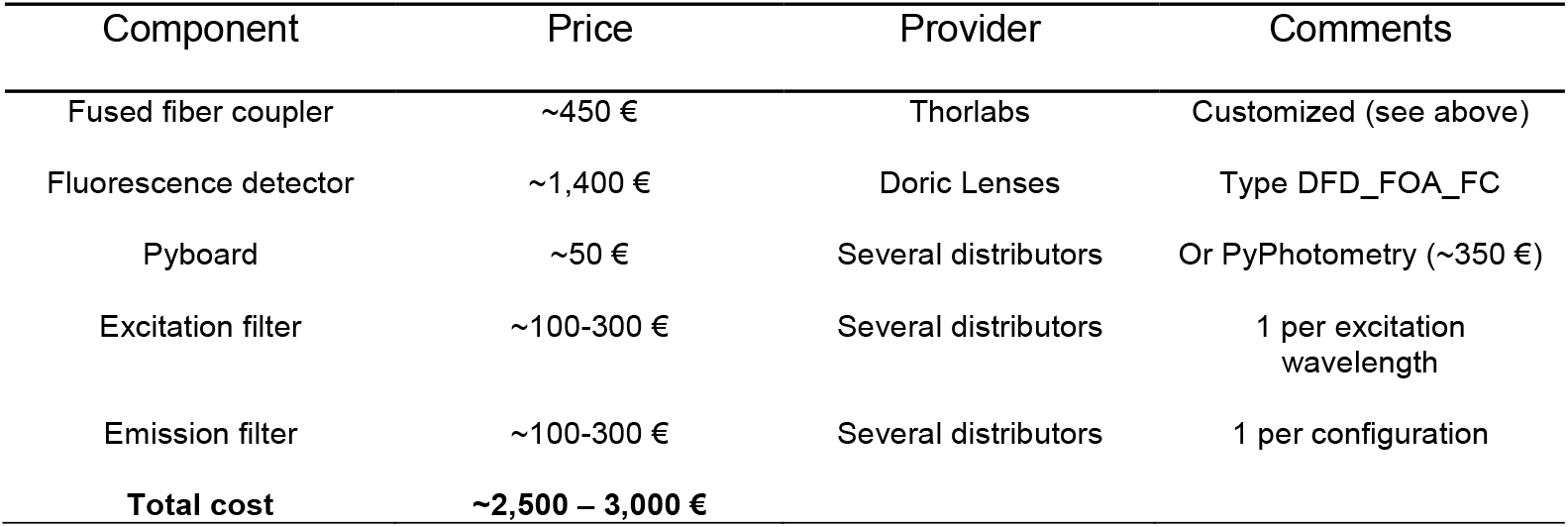
Components for the assembly of a fused fiber photometry system. Thanks to R&D efforts of many groups, the costs of fiber photometry systems dropped quite dramatically in recent years. Nonetheless, the FFP system proposed in the present work uses a conceptually different approach, resulting in a reduced number of components and opening a possibility for even further reduction of the price. Our approach is especially beneficial if a multicolor LED light source is already available in the laboratory. Note that light sources are not included in this table.

**Supplementary table 3.**
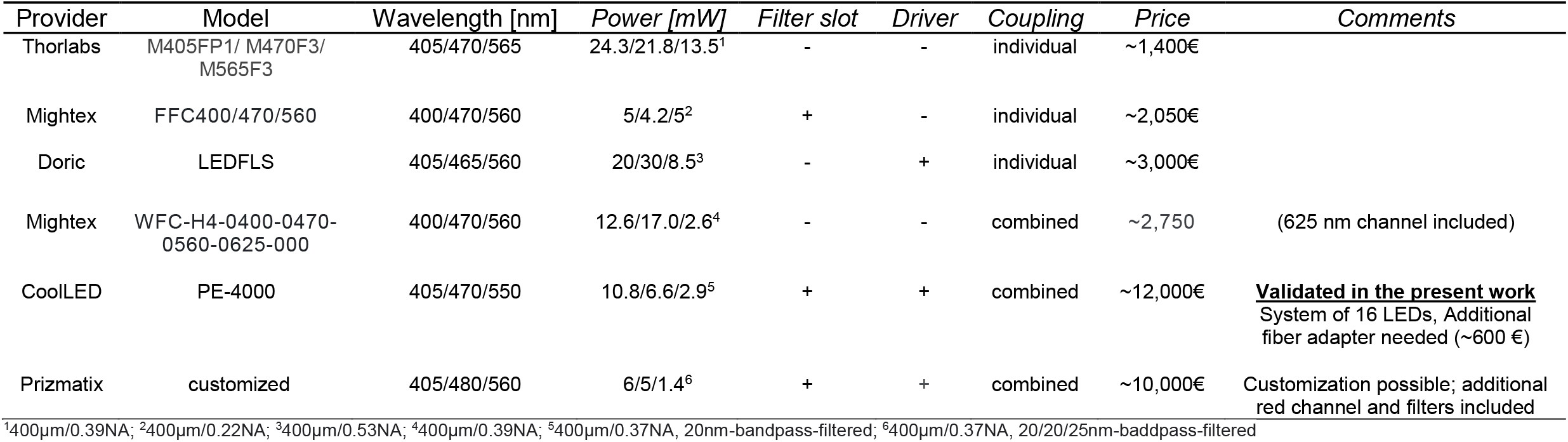
LED light sources for fiber photometry recordings. Note that power estimations for the CoolLED and Prizmatix systems are reported after optical filtering, while other light sources still need to be filtered. For LEDs without filter, external filter cubes + filters can be added (with increasing cost, and increasing coupling loss). For LEDs without driver, a separate LED driver is needed, e.g. the Cyclops LED driver by Open Ephys (∼500€) or pyPhotometry board (∼350€). This list provides selected examples of available light sources on the market most of which were not yet tested with the FFP system.

**Supplementary table 4.**
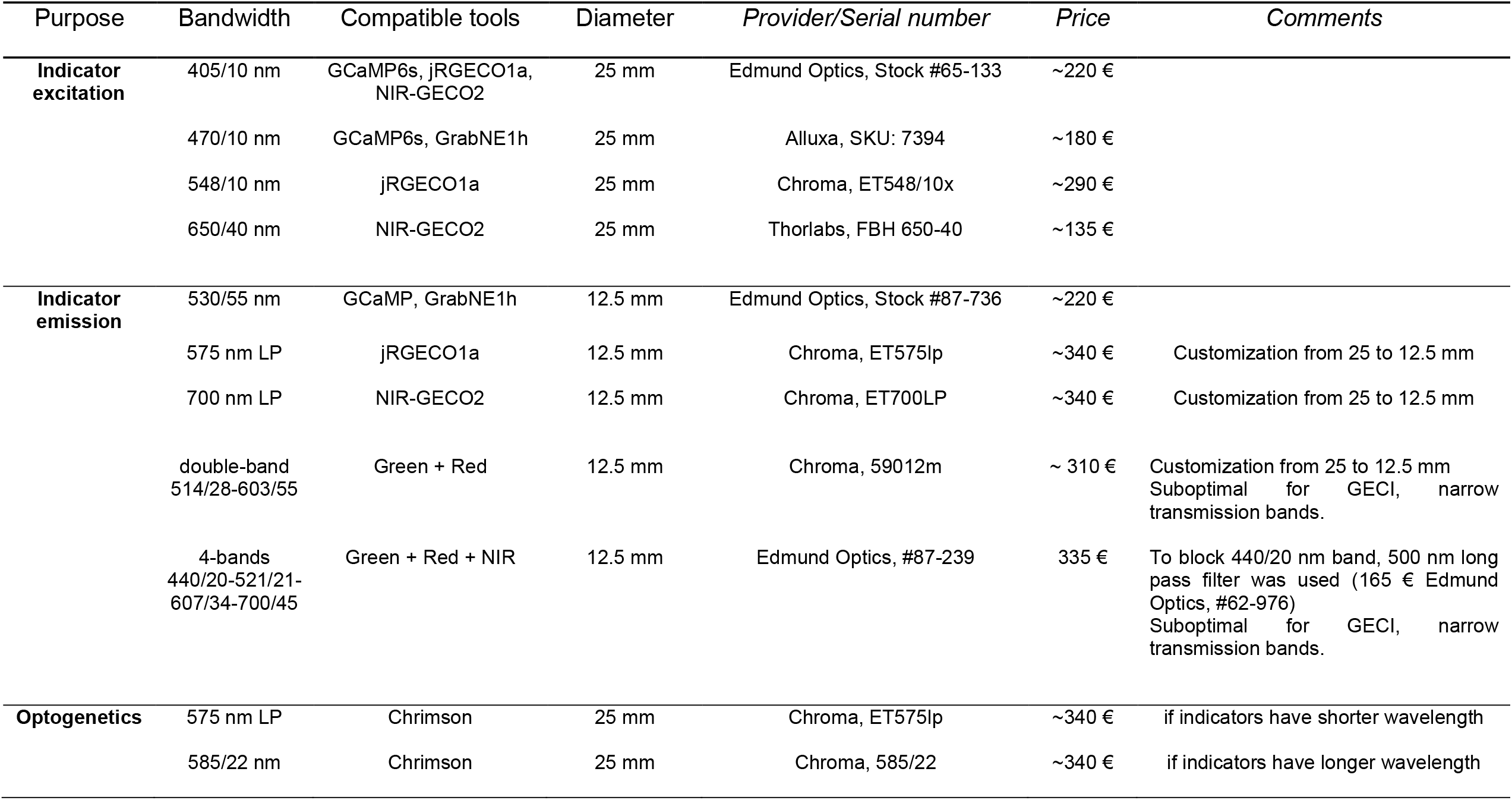
Filters used for FFP customization.

**Supplementary table 5.**
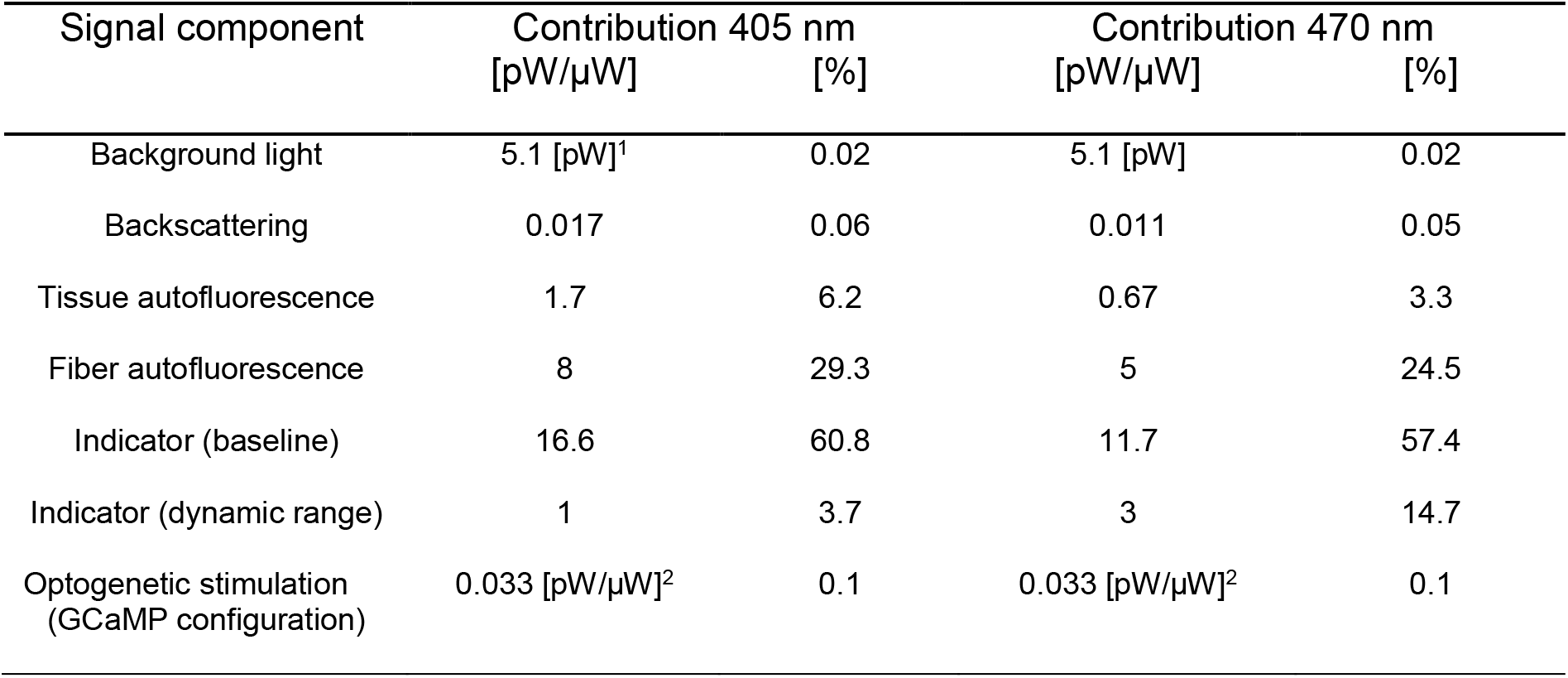
Relative contributions of fluorescence signal. Note that all values, except background light and optogenetic stimulation, are proportional to the power of excitation light, and thus reported in pW/µW. (1) Background light is a constant contribution and reported in pW, and (2) optogenetic stimulation is proportional to activation light, instead. Relative signal contributions are measured at excitation and activation light of 100 µW.

**Supplementary table 6.** Animals and viral constructs used in this study. Note that all transgenes were expressed in a non-conditional manner and therefore independent of the genetic background of the animals. Different mouse lines were used to minimize breeding excess in our mouse colonies, following the principle of the 3Rs (https://nc3rs.org.uk/who-we-are/3rs).

| Mouse ID | Line (genotype) | Hemisphere | Construct | Titer | Figure |
| --- | --- | --- | --- | --- | --- |
| 671 | GAD2 (cre+) | Right | rAAV2/9-CamKII-GCaMP6s | $1.25 \times 10^{13}$ | 1D-G; 1I,J; S6C,D |
| 751 | GAD2 (cre+) | Right | rAAV2/9 hSyn-jRGECO1a | $4.0 \times 10^{13}$ | 1L,M |
| 713 | GAD2 (cre+) | Right | rAAV2/9-CamKII-GCaMP6s | $1.25 \times 10^{13}$ | 1D-G; S6C,D |
| 11 | DAT-IRES-cre (cre-) | Both | rAAV2/9 CAG-NIR-GECO2 | $1.0 \times 10^{13}$ | S4B,C |
| 12 | DAT-IRES-cre (cre-) | Both | rAAV2/9 CAG-NIR-GECO2 | $5.0 \times 10^{12}$ | S4B-D |
| 92 | DAT-IRES-cre (cre+) | Right | rAAV2/9 CAG-NIR-GECO2 | $1.0 \times 10^{12}$ | 3B,D,E; S4B,C |
| 93 | DAT-IRES-cre (cre+) | Right | rAAV2/9 CAG-NIR-GECO2 +<br>rAAV2/9-hSyn-ChrimsonR-mCerulean (K176R) | $0.5 \times 10^{12}$<br>$1.5 \times 10^{12}$ | 3C-E; S4B/C; S5 |
| 15242 | C57BL/6J (NA) | Right | - | - | 3D,E; S4B,C; S6C,D |
| 15252 | C57BL/6J (NA) | Right | - | - | 3D,E; S4B,C; S6C,D |
| 15436 | C57BL/6J (NA) | Right | - | - | 3D,E; S4B,C; S6C,D |
| 15137 | C57BL/6J (NA) | Right | rAAV2/9-CamKII-GCaMP6s +<br>rAAV2/10 - hSyn-ChrimsonR-tdTomato (K176R) | $1.25 \times 10^{13}$<br>$1.13 \times 10^{13}$ | 1D-G; 2D-H; S6C,D |

**Supplementary figure 1.**
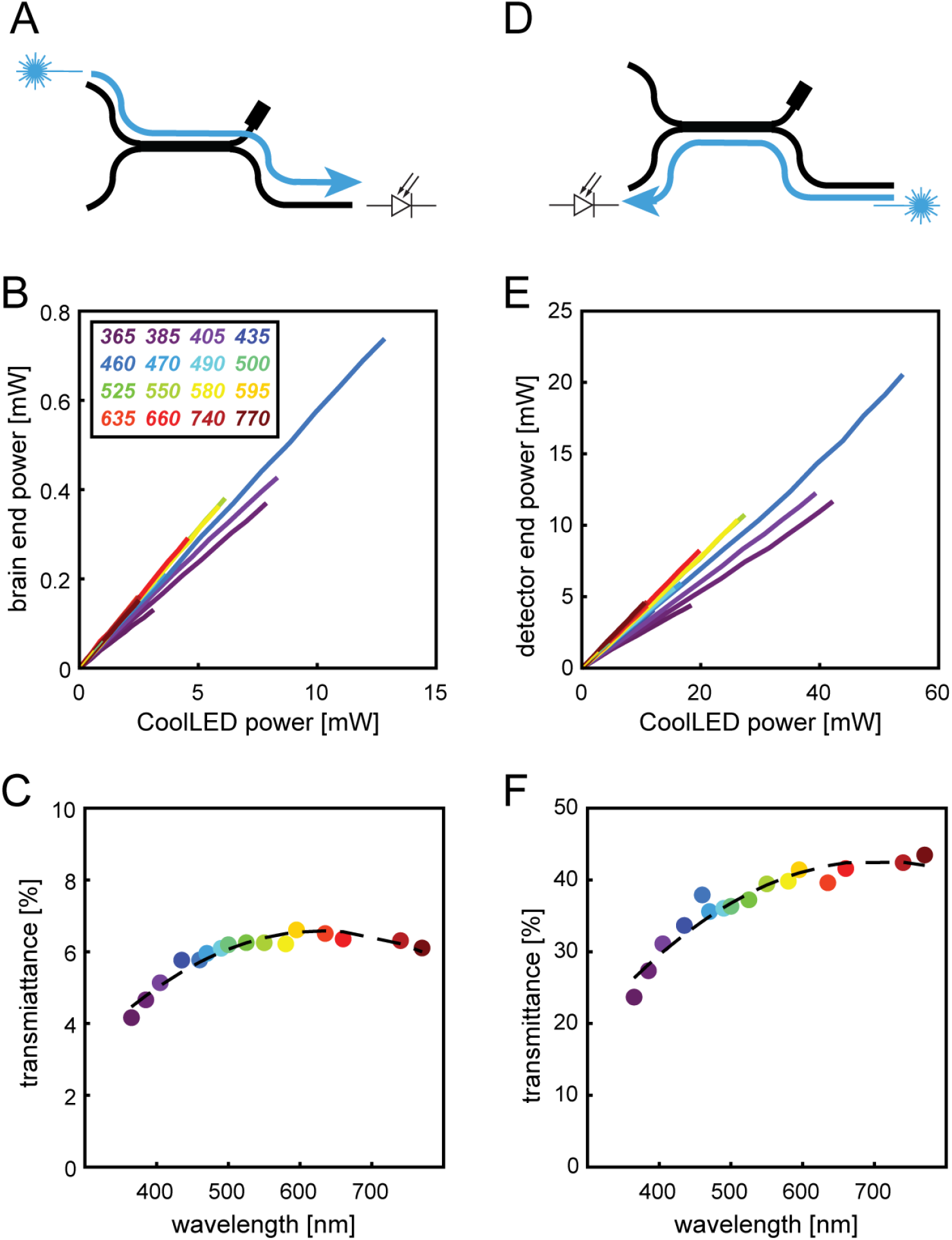
Optical properties of a fused fiber coupler. **(A)** Illustration of the measurement procedure to characterize the transmission efficiency of excitation (blue arrow), which propagates from the excitation port to the brain port though the fused fiber coupler. **(B)** Power at the brain port as a function of input light power from the LED light source for indicated wavelengths (in nm). **(C)** Transmittance of excitation light through the coupler defined as the ratio between power at the brain port and input power injected into the excitation port by a LED light source. Colors of circles match the wavelengths indicated in (B). **(D)** Illustration of the measurement procedure to characterize the transmission efficiency of emitted light (blue arrow), which propagates from the brain port to the detector port through the fused fiber coupler. **(E)** Power at the detector port as a function of input light power into the brain port. **(F)** Transmittance of emission light through the coupler defined as ratio between power at the detector port and input power into brain port.

**Supplementary figure 2.**
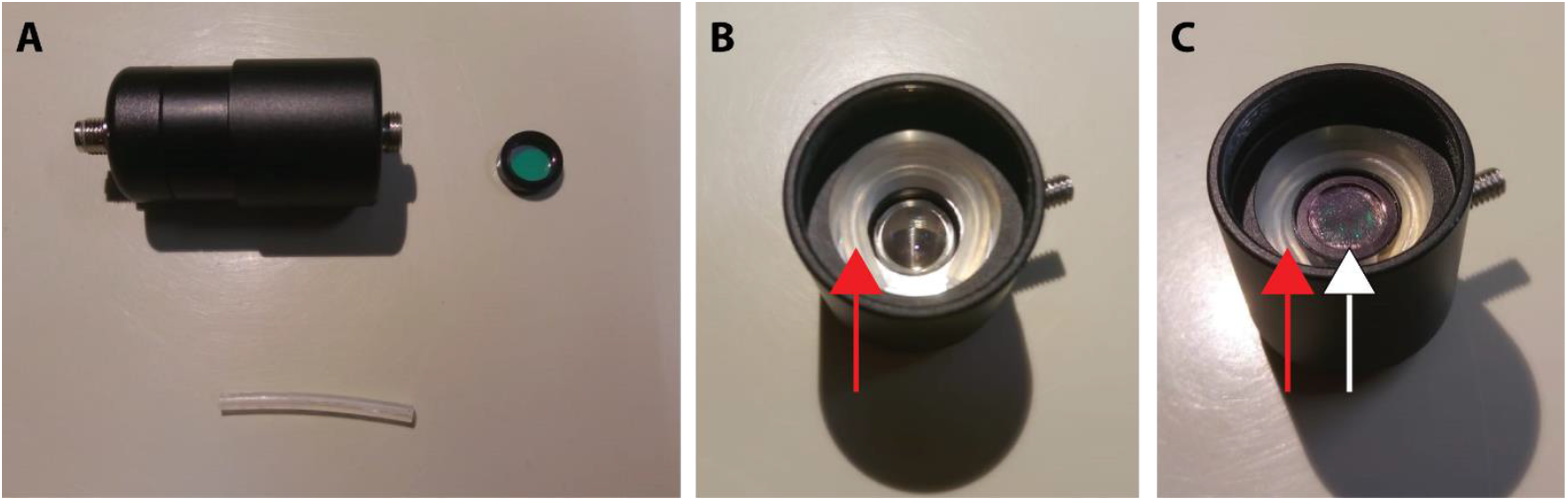
Filter installation in the photodetector. The 12.5 mm diameter filter was centered in the photodetector using a 3 mm diameter rubber tube. **(A)** Photodetector (closed), filter, and rubber tube. **(B)** Opened photodetector with the rubber tube (red arrow) in place to fix the filter. **(C)** Final installation of filter (white arrow), fixed in the photodetector by the rubber tube (red arrow).

**Supplementary figure 3.**
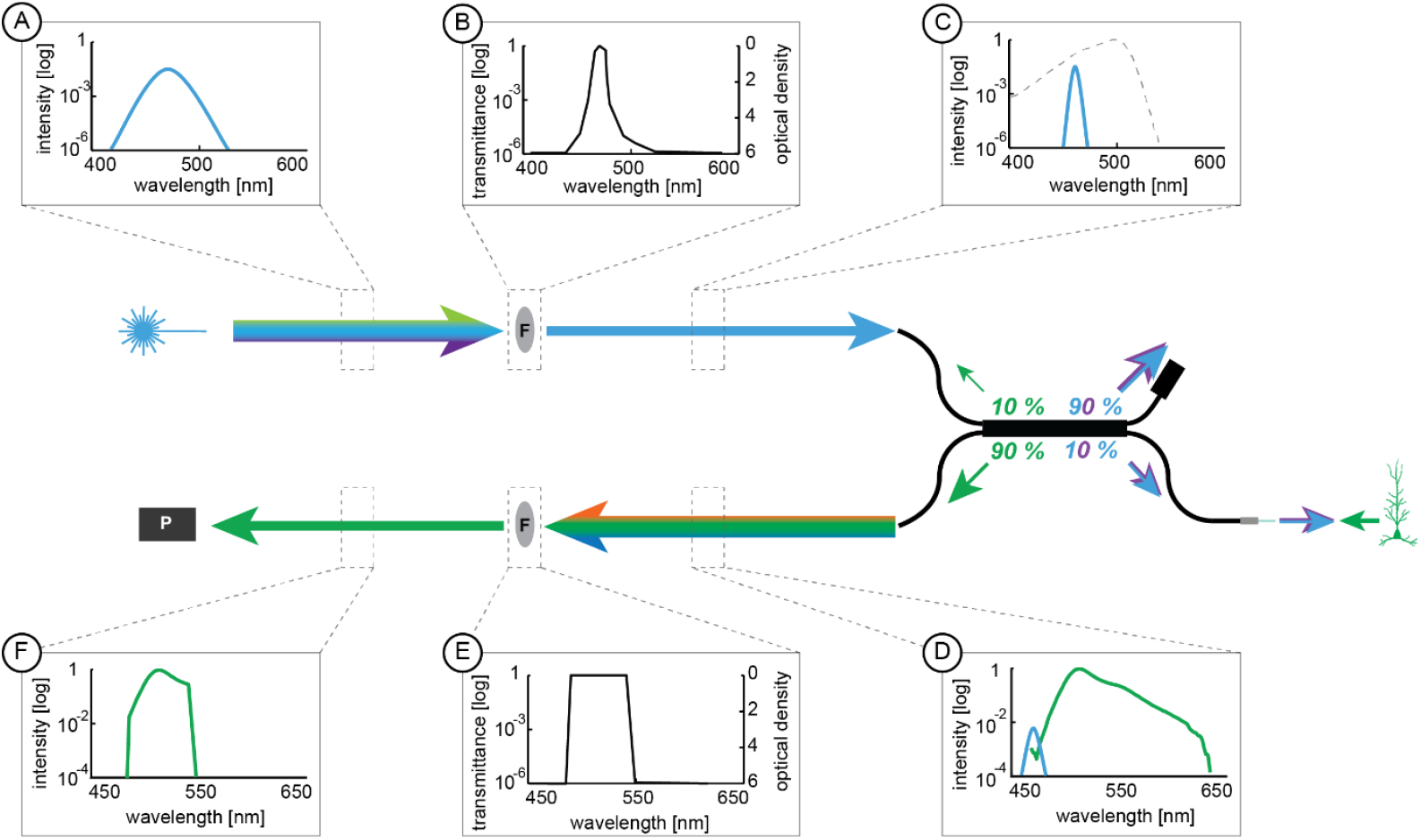
Spectral separation for excitation and emission light in FFP. The figure illustrates the propagation of excitation light from the light source to the brain (A – C) and emission light from the brain to the detector (D – F). The transformation of the spectra inside the FFC can be described as follows: The broad excitation light spectrum originating from the LED **(A)** is cleaned up with a band-pass filter **(B)** to restrict the excitation light to a narrower spectrum **(C)**, matching the peak excitation spectrum of the indicator fluorophore, but reduces optical contamination of the signal channel in the emission band. Dashed grey line: excitation spectrum of GCaMP6s. The indicator fluorophore emits light of longer wavelengths **(D)**, which is collected by the optical fiber at the brain end and propagates through the FFC to the detector. Before reaching the detector, the emission light is cleaned up with an additional band-pass filter to eliminate remaining excitation light and any additional optical noise (potentially originating from ambient light etc.) **(E)**. Finally, the filtered spectrum **(F)** is collected by the photodetector (P). Overall modulation of the detected light is strongly coupled to the modulation of light emission by the fluorescent indicator.

**Supplementary figure 4.**
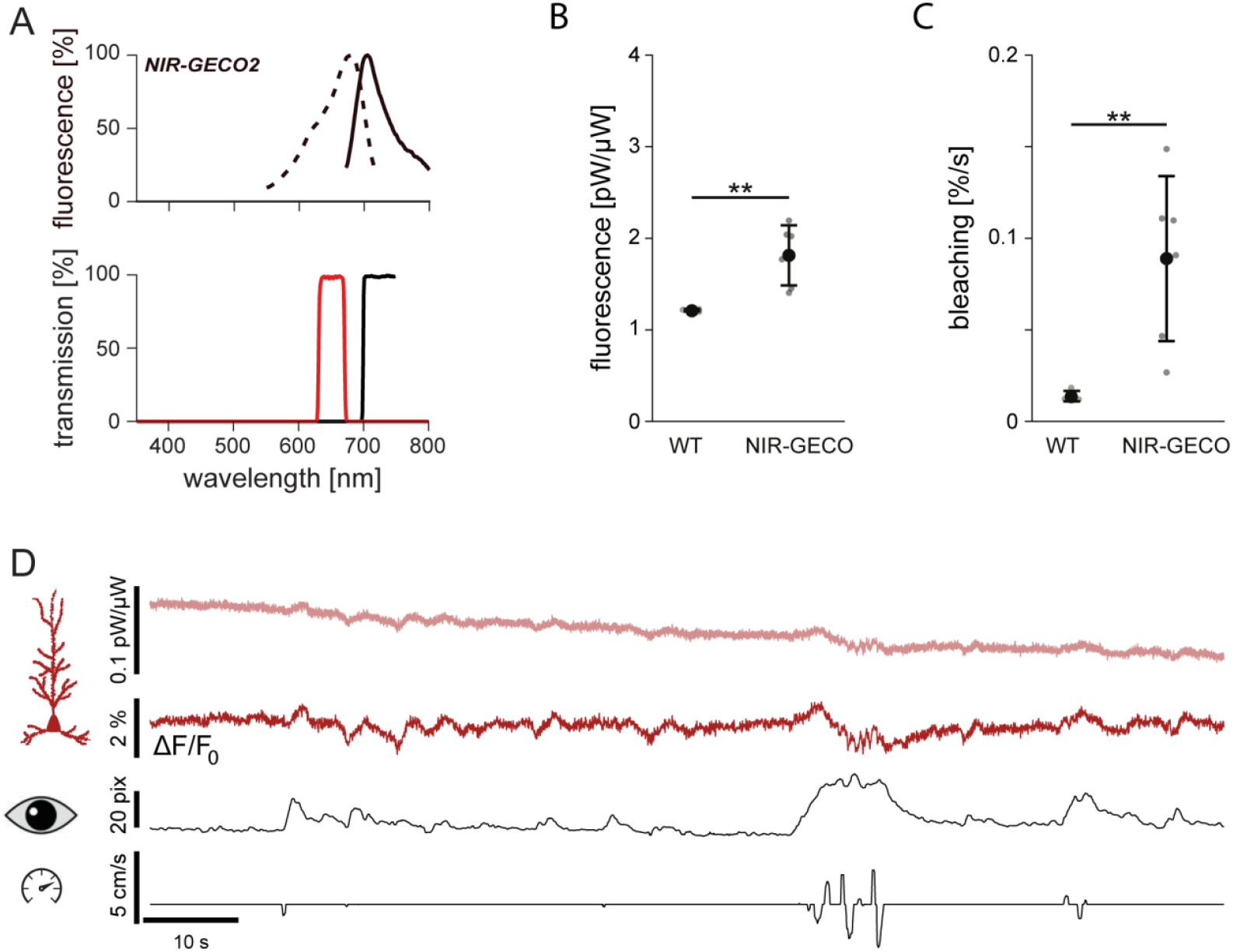
NIR-GECO2 recording using FFP. **(A)** Excitation and emission spectrum of NIR-GECO2 (top) and filter properties of excitation (650/40 nm, red) and emission filters (700 nm long pass, black; bottom) to record fluorescence from NIR-GECO2. **(B-C)** Mean ± standard deviation of the absolute fluorescence (*p* = 0.007, Welch’s two-sample t-test; **B**) and bleaching (*p* = 0.008, Welch’s two-sample t-test; **C**) in NIR-GECO2-injected mice (n = 6 hippocampi in N = 4 mice) and non-injected controls (n = 3 hippocampi in N = 3 mice). **(D)** Representative raw data traces of 2 min duration: raw traces for 650 nm excitation 650 nm-baseline-fluorescence corrected calcium signal (ΔF/F_0_), pupil size, and animal locomotion (from top to bottom). **: p < 0.01.

**Supplementary figure 5.**
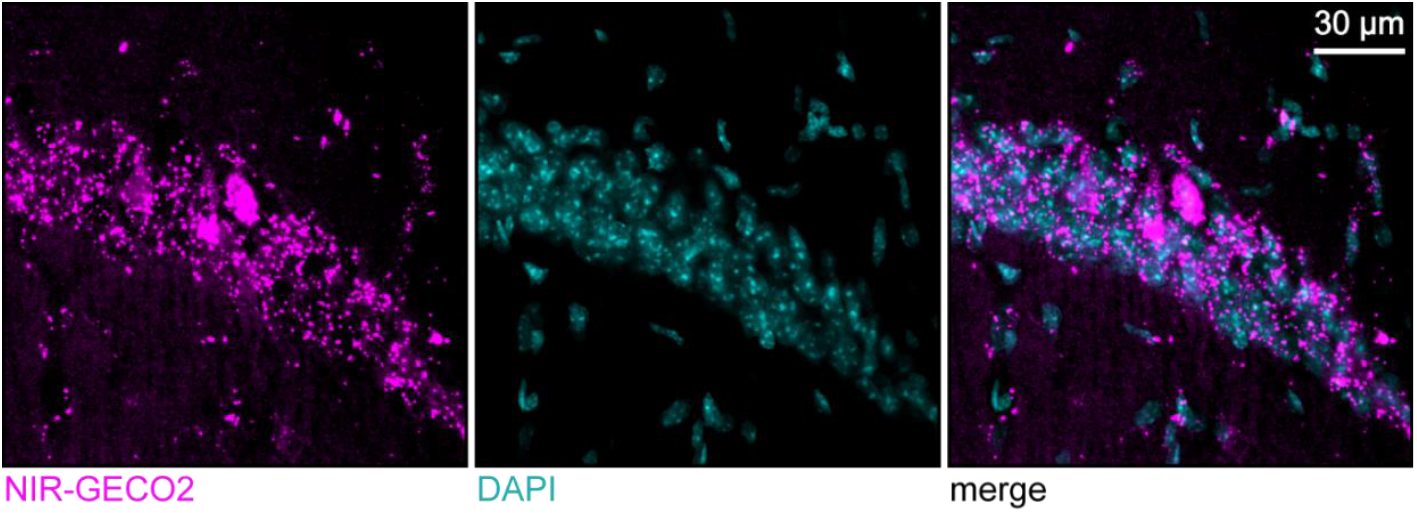
Protein aggregates of NIR-GECO2. NIR-GECO2 expressing (magenta) hippocampal neurons in stratum pyramidale of area CA1 (visualized by DAPI-staining, cyan). Note accumulation of potentially unfunctional NIR-GECO2 protein aggregates, evident by the bright puncta (magenta).

**Supplementary figure 6.**
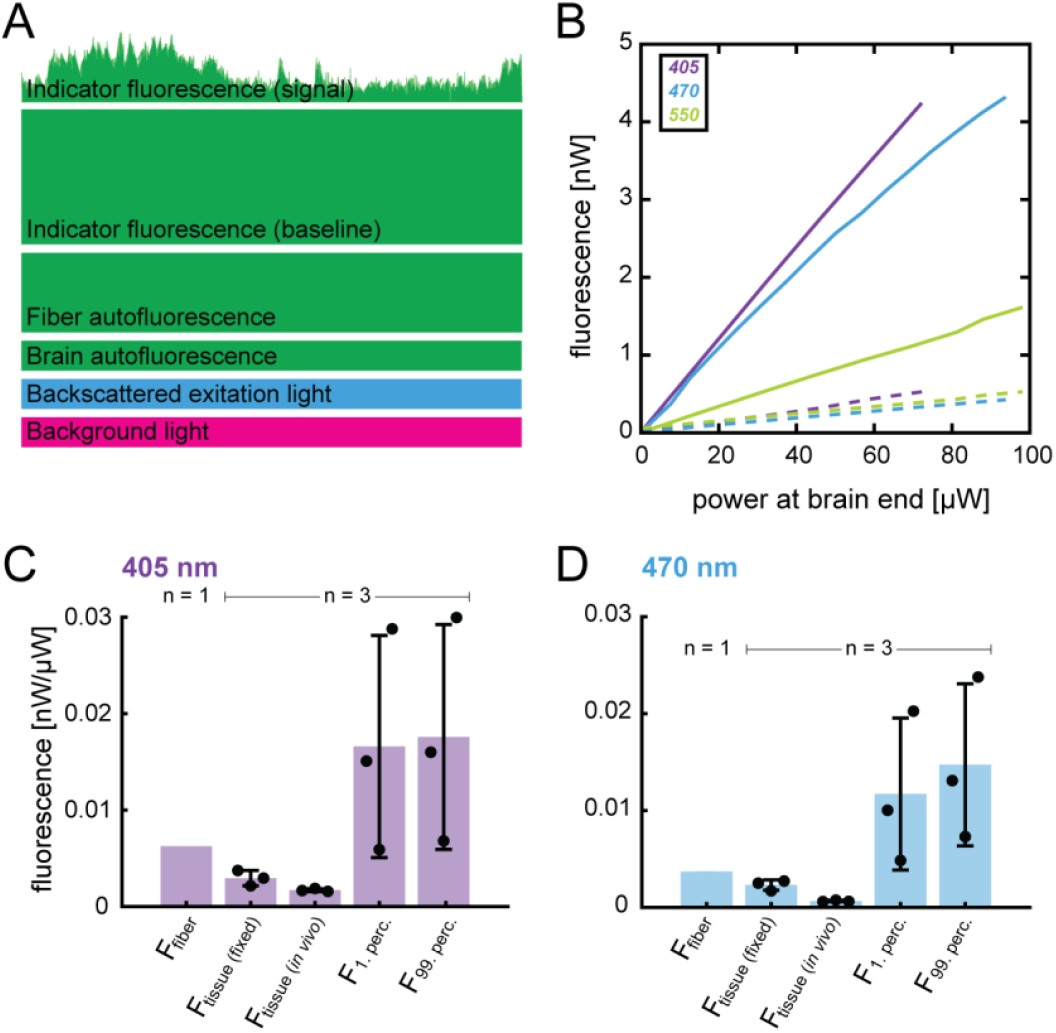
Conceptual model of the fluorescence signal and empirical assessment of contributing fluorescence sources. **(A)** Illustration of the composition of the detected fluorescence signal. **(B)** Autofluorescence of the fiber measured by the photodetector as a function of excitation power at the brain port before (solid line) and after 12-hours of photobleaching (dashed line). **(C, D)** Signal contributions for excitation of GCaMP6s with light of 405 (C) and 470 nm (D): Fluorescence (F) of the optical fiber, brain tissue (fixed or *in vivo*) and upper and bottom percentile of the calcium transients.

## Footnotes

1 A femtowatt photoreceiver can still be used with ratiometric indicators but the possibility to detect both spectral contributions from CFP and YFP independently is lost in this case as the detector cannot distinguish the emission spectra.

